# Bacterial Shedu immune nucleases share a common enzymatic core regulated by diverse sensor domains

**DOI:** 10.1101/2023.08.10.552793

**Authors:** Yajie Gu, Huan Li, Amar Deep, Eray Enustun, Dapeng Zhang, Kevin D. Corbett

**Author notes:** Correspondence (D.Z.); (K.D.C.).

## Abstract

Prokaryotes encode diverse anti-bacteriophage immune systems, including the single-protein Shedu nuclease. Here we reveal the structural basis for activation of *Bacillus cereus* Shedu. In the inactive homotetramer, a key catalytic residue in Shedu’s nuclease domain is sequestered away from the catalytic site. Activation involves a conformational change that completes the active site and promotes assembly of a homo-octamer for coordinated double-strand DNA cleavage. Removal of Shedu’s N-terminal domain ectopically activates the enzyme, suggesting that this domain allosterically inhibits Shedu in the absence of infection. Bioinformatic analysis of nearly 8,000 Shedu homologs reveals remarkable diversity in their N-terminal regulatory domains: we identify 79 domain families falling into eight functional classes, including diverse nucleic acid binding, enzymatic, and other domains. Together, these data reveal Shedu as a broad family of immune nucleases with a common nuclease core regulated by diverse N-terminal domains that likely respond to a range of infection-related signals.

## Introduction

The ongoing evolutionary conflict between bacteria and their viral pathogens (bacteriophages or phages) has driven the development of myriad bacterial immune systems that fight infection, and an equally complex array of viral factors that counter these systems ^1,2^. The most well understood bacterial immune systems, restriction-modification systems and CRISPR-Cas systems, are found in a majority of bacterial genomes ^3^. Recent advances in understanding of bacterial genome organization, combined with the availability of tens of thousands of bacterial genome sequences, have enabled the discovery of dozens of novel immune systems that are more sparsely distributed across bacterial populations ^3–8^.

Strikingly, many bacterial immune systems are now understood to be evolutionary predecessors of eukaryotic innate-immune systems, providing a new understanding of how innate immunity has evolved across life ^9–18^.

Bacterial immune systems use a startling variety of mechanisms to sense phage infection and trigger a defensive response. Many immune system effectors are nucleases that specifically target foreign DNA, such as restriction enzymes that target DNA lacking specific modifications ^19,20^, or CRISPR-Cas systems that recognize and cleave specific DNA sequences ^21–24^. These systems typically eliminate infections without significant collateral damage to the host’s genome. Many other effector nucleases, including the CBASS-associated nucleases NucC and Cap4, are non-specific and cause catastrophic damage to both the phage and host genomes once activated by an infection signal. Activation of these non-specific nuclease effectors can lead to cell death, in a so-called abortive infection mechanism ^25,26^.

The Shedu system is among the simplest known anti-phage immune systems, comprising a single protein (SduA; here termed Shedu). Shedu is found in 2.3% of bacterial genomes, often within “defense islands” enriched for anti-phage and other defense systems ^5^. The Shedu protein is proposed to act as a nuclease, with a conserved DUF4263 (Pfam ID: PF14082) domain that belongs to the PD-(D/E)XK nuclease superfamily ^5,27^. Specific modification of phage DNA can suppress Shedu-mediated anti-phage immunity and *in vitro* nuclease activity ^27^. Still unknown, however, is how Shedu’s nuclease activity is regulated in response to phage infection.

Here, we use cryoelectron microscopy (cryoEM) to determine two structures of Shedu from *Bacillus cereus* strain B4264, revealing an oligomerization-associated conformational switch between inactive and active nuclease states. We find that this conformational switch is regulated by the protein’s N-terminal domain, which adopts a structure related to the nucleic acid-binding sPC4/Whirly fold. Through an exhaustive examination of nearly 8,000 Shedu homologs throughout prokaryotes, we identify 79 distinct N-terminal domain families fused to the common C-terminal nuclease domain. These domains enable classification of Shedu homologs into eight broad families, each of which likely senses a distinct signal associated with phage infection or other stimuli. Overall, our data reveal Shedu as a diverse family of immune nucleases, with a common nuclease core that is allosterically controlled by diverse N-terminal domains.

## Results

### *B. cereus* Shedu is a non-abortive immune nuclease

The single-gene Shedu immune system from *Bacillus cereus* B4264 has previously been shown to protect a *Bacillus subtilis* host from infection by two closely related phages, rho14 and phi105 ^5^. To test whether Shedu’s anti-phage activity is dependent on its predicted PD-(D/E)XK nuclease activity, we tested mutants of two highly conserved active site residues within this family of nucleases: motif II residue D249, and motif III residue E264. We found that mutation of either D249 or E264 to alanine eliminated Shedu’s ability to protect against both rho14 and phi105 (**Figure 1A**). To determine whether *B. cereus* Shedu is an abortive infection system, we infected *B. subtilis* cells expressing Shedu or the E264A mutant (SduA gene integrated into the bacterial genome) with phages rho14 or phi105 in liquid culture. In contrast to known abortive infection systems, cells expressing Shedu did not die in a high-multiplicity infection (MOI 10; **Figure 1B-C, Figure S1**). Cells expressing the Shedu E264A mutant were efficiently infected, further demonstrating that the protein’s predicted nuclease activity is responsible for its immune function. These data suggest that *B. cereus* Shedu specifically targets incoming phage DNA for destruction, rather than nonspecifically cleaving host and phage DNA upon activation as in abortive infection systems.

**Figure 1.**
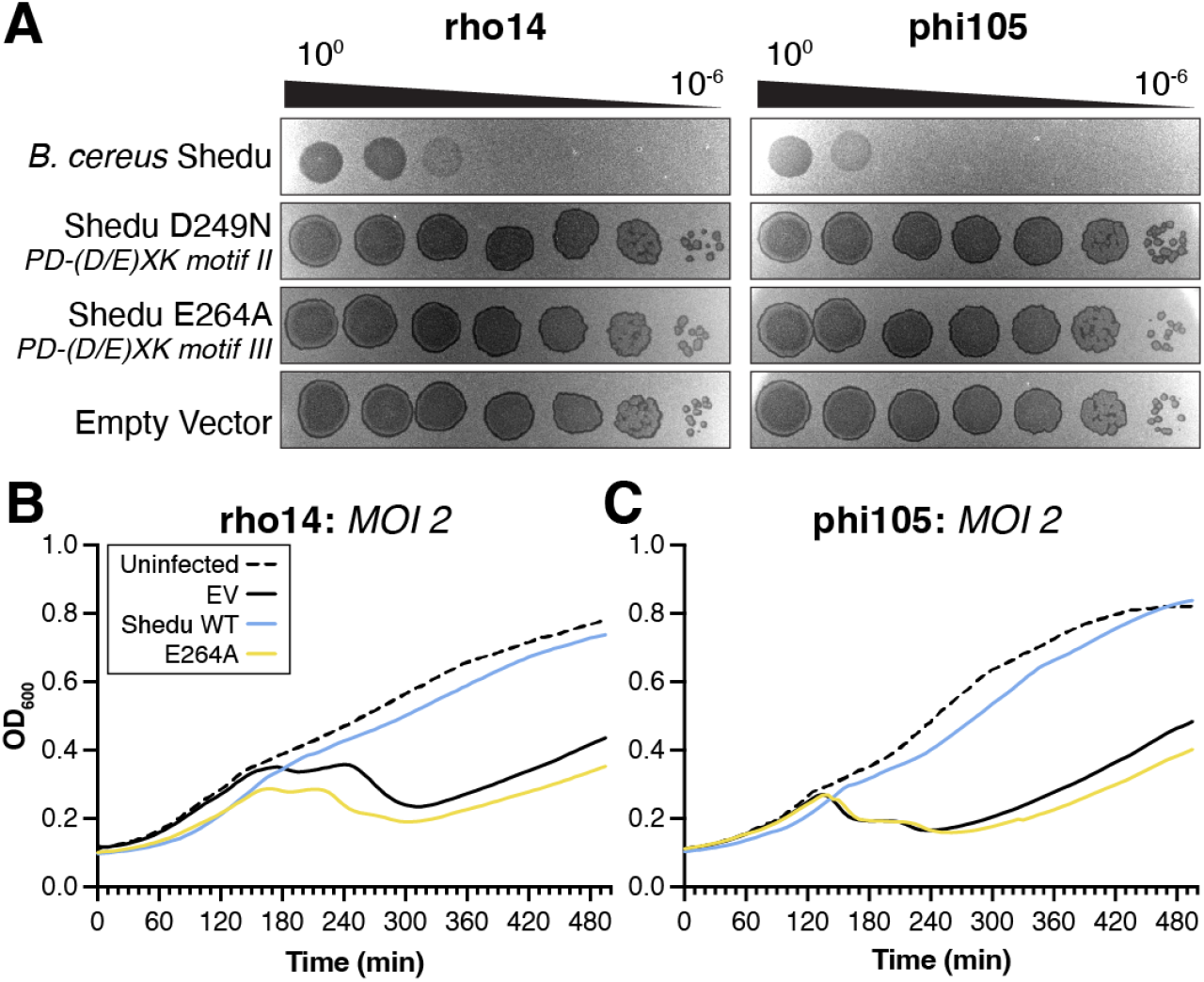
*B. cereus* Shedu is a non-abortive immune system. (A) Plaque assays with *B. subtilis* cells with genome-integrated *B. cereus* B4264 Shedu: wild-type, D249N motif II mutant, or E264A motif III mutant. Phages used were rho14 (left) and phi105 (right), with six ten-fold dilutions from a high-titer working stock. Images are representative of three independent trials. (B) and (C) Growth curves of *B. subtilis* cells with *B. cereus* B4264 Shedu integrated into their genome (empty vector: black lines, wild-type: blue lines, E264A: yellow lines) with rho14 (panel (B)) or phi105 (panel (C)) at a multiplicity of infection (MOI) of 2 at time zero. See **Supplemental Figure S1** for growth curves at other MOIs. Lines represent average values from triplicate measurements; see **Supplemental Figure S1** for error bars. Both phages are temperate, so cultures recover even at high MOI due to integration of the phage into the host genome.

### *B.* cereus Shedu adopts an inactive tetramer conformation

We next purified full-length *B. cereus* Shedu for biochemical and structural analysis. Sequence alignments show that the protein possesses three domains: an N-terminal domain of unknown architecture, a short central linker domain, and a C-terminal predicted nuclease domain (**Figure 2A**). Using size exclusion chromatography coupled to multi-angle light scattering (SEC-MALS), we found that *B. cereus* Shedu exists predominantly as a homotetramer in solution, with a small fraction of a larger oligomer consistent with a hexamer or octamer species (**Figure S2A**). We performed negative-stain electron microscopy on purified *B. cereus* Shedu, and observed reproducible elongated, three-lobed particles averaging 21 nm in length (**Figure 2B-C**). We next used cryoelectron microscopy (cryoEM) to determine a 2.91 Å resolution structure of *B. cereus* Shedu (**Supplemental Figure 2, Supplemental Table S1**). In agreement with our SEC-MALS data, the structure reveals an extended architecture with four protomers’ nuclease domains forming a central homotetramer, and the N-terminal domains extending outward from each end of the complex. In the structure, the N-terminal domains are disordered, and the central linker domains are partially disordered, due to flexibility within the complex (**Figure 2D-E**). The Shedu nuclease domain adopts an elaborated PD-(D/E)XK nuclease fold, with the closest structural homolog being *Thermococcus kodakarensis* EndoMS ^28^, a restriction endonuclease-related protein that specifically cleaves mismatched DNA (**Figure 2F**). Shedu possesses all active site motifs of the PD-(D/E)XK nuclease family, including motif I (E204 in *B. cereus* Shedu), motif II (D249), motif III (E264 and K266), and motif IV (Q295). These residues are all clustered on the surface of the Shedu protomer, with the exception of D249. This residue is located on a long loop that we term the motif II loop and is positioned more than 20 Å away from other active site residues (**Figure 2F**). The motif II loop is stably docked against a long α-helix (αC) through hydrophobic interactions of V248 and Y252 in the motif II loop with Y307 and I310 of the αC helix (**Figure 2F**). Given that motif II is highly conserved and required for Shedu’s anti-phage activity, the location of the motif II loop suggests that our structure represents an inactive, autoinhibited state.

**Figure 2.**
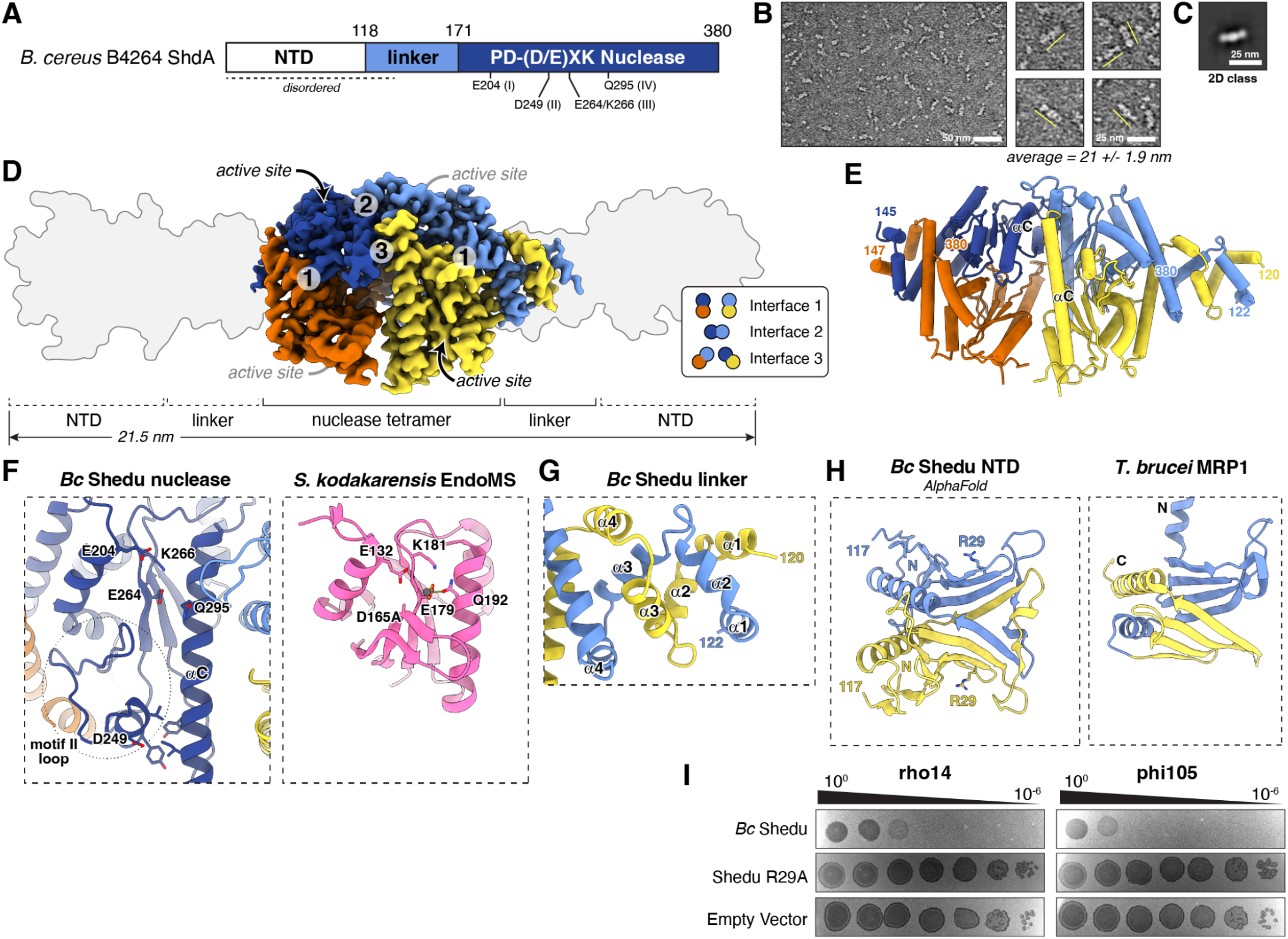
Structure of an inactive Shedu homotetramer. (A) Domain structure of *Bacillus cereus* B4264 Shedu, with N-terminal domain (NTD) white, linker domain light blue, and C-terminal predicted PD-(D/E)XK Nuclease (DUF4263) domain dark blue with conserved nuclease motifs noted. (B) Negative stain electron micrographs of purified full-length *B. cereus* Shedu. *Left:* Representative image (scale bar: 50 nm). *Right:* Individual particle images (highlighted with yellow lines; scale bar: 25 nm). Average length of 60 particles manually measured across 7 images: 21 +/-1.9 nm. (C) 2D class average from negative-stain electron microscopy of *B. cereus*. Scale bar: 25 nm. (D) CryoEM density for a *B. cereus* Shedu homotetramer (see **Supplemental Figure S2A** for SEC-MALS analysis), with the four protomers colored dark blue, light blue, orange, and yellow. Approximate locations of each active site are indicated (for orange and light blue subunits, active sites are not visible). The NTD and linker domains are largely disordered, and their inferred locations are indicated by the gray outline (produced from a composite model including an AlphaFold prediction of the NTD structure; see panel (F) and **Supplemental Figure S2B-C**). Numbers 1-3 indicate the locations of the three dimer interfaces that make up the tetramer. Inset indicates the pairs of subunits that are linked by each dimer interface. See **Supplemental Figure S2D-H** for structure determination workflow. (E) Cartoon view of the *B. cereus* Shedu homotetramer, colored as in panel (B) (F) *Left:* Closeup of the Shedu active site, with residues comprising motifs I-IV shown in sticks and labeled. The extended motif II loop is circled, with hydrophobic residues docking it to the extended ɑC helix shown as sticks. *Right*: Closeup of the *T. kodakarensis* EndoMS active site (PDB ID 5GKE)^28^, with residues comprising motifs I-IV shown in sticks and labeled. (G) Closeup of the Shedu linker domain dimer, with the two protomers colored light blue and yellow. The helical spiral interface is an extension of dimer interface 1. This domain is ordered in one half of the complex, and disordered in the other (see panel (B)). (H) *Left:* AlphaFold model of a dimer of the Shedu NTD, with protomers colored light blue and yellow. Conserved basic residue R29, located on the surface equivalent to the RNA-binding face of MRP1, is shown as sticks. See **Supplemental Figure S2B-C** for details of AlphaFold modeling. *Right:* Structure of a *T. brucei* MRP1 monomer (PDB 2GIA)^29^, with N-terminal subdomain colored light blue and C-terminal subdomain colored yellow. (I) Plaque assays with *B. subtilis* cells with *B. cereus* B4264 Shedu integrated into their genome: wild-type or R29A mutant. Phages used were rho14 (left) and phi105 (right), with six ten-fold dilutions from a high-titer working stock. Images are representative of three independent trials. Images for wild-type Shedu and Empty Vector images are the same as in Figure 1A.

The Shedu homotetramer is assembled from four PD-(D/E)XK nuclease domains, each of which interacts with other subunits through three distinct interfaces that we term interfaces 1, 2, and 3 (**Figure 2D**). Interface 1 is the largest at 1700 Å^2^ of surface area per protomer, and links the pairs of protomers that define each end of the extended complex. The two nuclease domain dimers defined by interface 1 are near-identical to one another, with an overall Cα r.m.s.d. across both protomers of ∼0.2 Å. Interface 2 resembles the canonical dimer interface of restriction endonucleases, including *T. kodakarensis* EndoMS. In the Shedu tetramer structure, one pair of protomers (colored dark blue and light blue in **Figure 2E**) is tightly associated across this interface, burying 1,250 Å^2^ of surface area per protomer.

The opposite pair of protomers (colored orange and yellow in **Figure 2E**) is rotated away from one another and bury only 190 Å^2^ per protomer. This difference lends a distinct asymmetry to the overall Shedu tetramer structure. Finally, the smaller interface 3 (475 Å^2^ per protomer) links the long αC helix of subunits positioned diagonally across interface 2. This interface is asymmetric, with R305, Q308, N309, and Y312 of one protomer packing against Y312, Y315, and F323 of the opposite protomer.

While the Shedu homotetramer contains four nuclease active sites, these sites are not positioned for coordinated cleavage of a double-stranded DNA. The closest pairs of active sites are those sites positioned across the tight interface 2, but these two sites are rotated away from one another by nearly 45° compared to an active nuclease homodimer like *T. kodakarensis* EndoMS. Thus, not only is each active site in an autoinhibited state with motif II residue D249 positioned far from other active site residues, the overall architecture of the Shedu tetramer does not appear to be compatible with double-strand DNA cleavage.

Within the overall Shedu homotetramer structure, each pair of subunits defined by interface 1 is also associated through their central linker domains (residues 118-170), which form a tight right-handed spiral comprising four short α-helices from each protomer, and extend outward from the nuclease domains (**Figure 2G**). The N-terminal domains (residues 1-117) are disordered in our structure, likely due to flexibility in the linker domains. We attempted to determine the structure of the isolated Shedu N-terminal domain, but were unable to express and purify this domain on its own. Instead, we used AlphaFold 2 to generate a confident structural prediction for this domain as a β-sheet rich fold similar in architecture to sPC4/Whirly proteins including the *T. brucei* MRP1 and MRP2 RNA binding proteins (**Supplemental Figure 2B-C**) ^29^. sPC4/Whirly proteins typically comprise a tandem repeat of β-β-β-β-α secondary structure elements, and the Shedu N-terminal domain model aligns with one of these repeats of MRP1. Modeling a homodimer of Shedu N-terminal domains in AlphaFold 2 results in a predicted structure displaying a dimer architecture very similar to a *T. brucei* MRP1 monomer, with the C-terminal β-strand and α-helix of each N-terminal domain showing a domain swap compared to MRP1 (**Figure 2H**). Thus, the Shedu homotetramer is assembled from two dimers, each of which is arranged as a parallel assembly of dimerized N-terminal sPC4/Whirly domains, linker domains, and DUF4263 nuclease domains. Given that many sPC4/Whirly fold proteins bind single-stranded nucleic acids, such as MRP1/MRP2 ^29^, PC4 ^30^, Whirly ^31^, and PurA ^32^, we propose that this domain may serve as an infection sensor by binding nucleic acids. In agreement with this idea, mutation of a highly conserved arginine residue (R29) on the domain’s putative nucleic acid binding surface eliminated Shedu’s anti-phage immunity (**Figure 2I**).

### The Shedu N-terminal domains regulate nuclease activity

Our structure of full-length *B. cereus* Shedu revealed a tetrameric assembly with apparently inactive nuclease domains. Consistent with this observation, overexpression of full-length wild-type *B. cereus* Shedu in *E. coli* is not toxic to cells (**Supplemental Figure S3A-D**). In contrast, a truncated construct lacking the N-terminal and linker domains (residues 171-380; Shedu-ΔNL) was highly toxic to *E. coli* (**Supplemental Figure S3A-D**). This toxicity depended on an intact nuclease active site, as the E264A mutant was not toxic. These data suggest that Shedu’s N-terminal and linker domains allosterically regulate the protein’s nuclease activity, maintaining the enzyme in an inactive state in the absence of infection.

To determine the mechanism of allosteric control by the Shedu N-terminal and linker domains, we purified the nuclease-dead E264A mutant of Shedu-ΔNL. By SEC-MALS, we found that Shedu-ΔNL adopts a stable octamer rather than the predominantly tetrameric form adopted by the full-length protein (**Supplemental Figure S3E**). We next determined the structure of the Shedu-ΔNL octamer by cryoEM to a resolution of 3.19 Å (**Supplemental Figure 4, Supplemental Table S1**). We also locally refined a single tetramer within the Shedu-ΔNL octamer to a resolution of 2.77 Å. The structure reveals a Shedu-ΔNL octamer with two tetramers associated side-by-side (**Figure 3A-B**). Compared to the full-length Shedu tetramer, each tetramer in the Shedu-ΔNL undergoes a significant conformational change in which the two subunits that were previously rotated away from one another across dimer interface 2 (colored orange and yellow in **Figure 3C**) move and rotate toward one another (**Figure 3C, Supplemental Movie S1**). The resulting close protomer-protomer packing across the newly-formed second interface 2 forces the long αC helix of two protomers associated across interface 3 (dark blue and yellow in **Figure 3C**) to kink by 90°. Kinking of the αC helices exposes interface 3 residues Y312 and Y315 in these two protomers, creating the octamer interface (**Figure 3B-C**).

**Figure 3.**
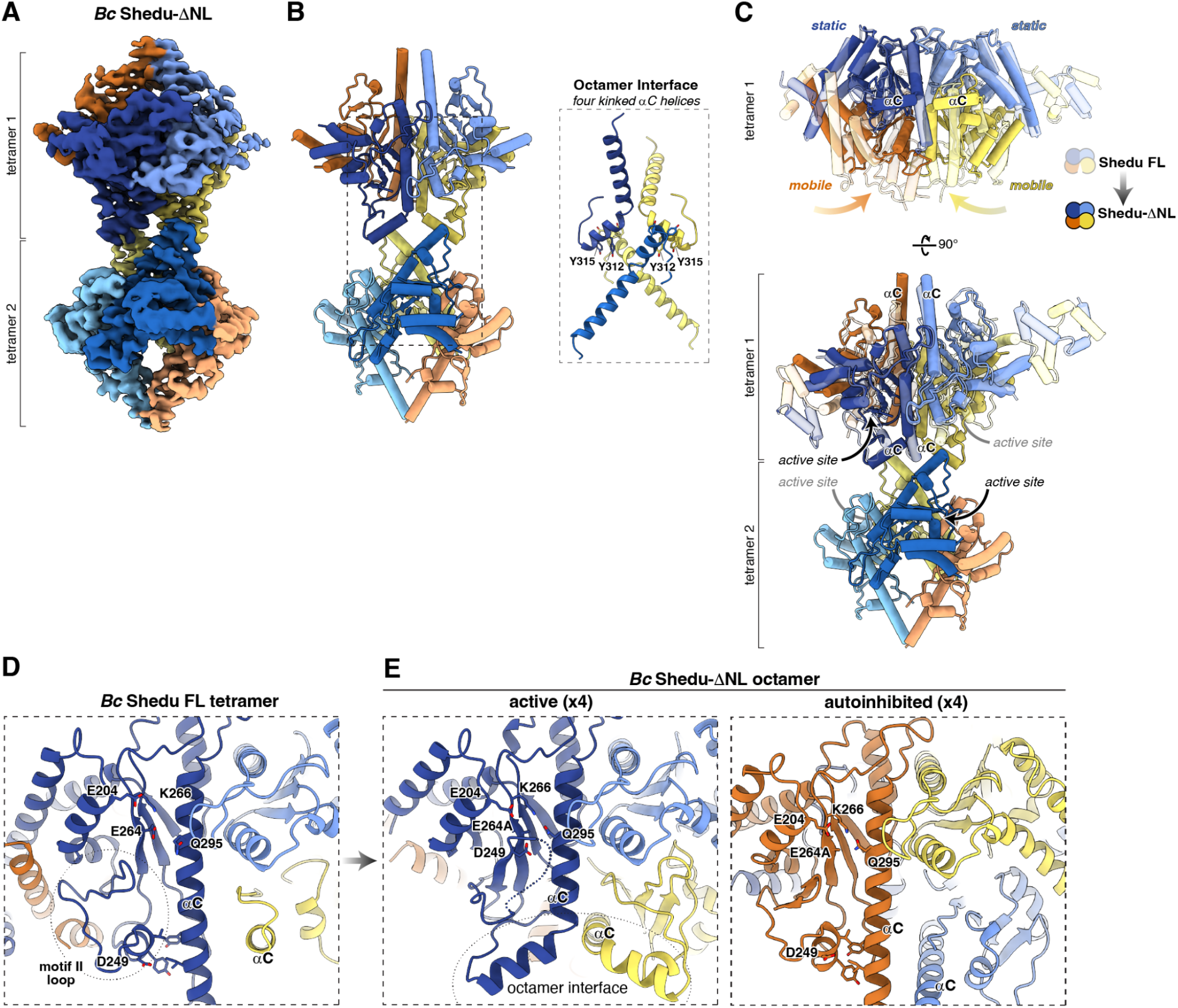
Structure of an active Shedu octamer. (A) CryoEM structure for the *B. cereus* Shedu-ΔNL octamer (see **Supplemental Figure S3E** for SEC-MALS analysis), with protomers colored as in Figure 2B. See **Supplemental Figure S4** for structure determination workflow. (B) Cartoon view of the *B. cereus* Shedu-ΔNL octamer, colored as in panel (B). Inset shows the four kinked ɑC helices that make up the octamer interface. Residues Y312 and Y315, which interact with one another through the octamer interface, are shown as sticks. (C) Structural overlay of the Shedu FL tetramer structure (semitransparent) with one tetramer in the Shedu-ΔNL (solid). The light blue and dark blue subunits (marked “static”) do not change position between the two states, while the yellow and orange subunits (marked “mobile”) rotate and shift toward one another. At the same time, the ɑC helices of the dark blue and yellow protomers kink to form the octamer interface (see rotated lower panel) and release the motif II loop to activate the enzyme. See **Supplemental Movie S1**. (D) Closeup of the *B. cereus* Shedu active site in the tetramer state, with residues comprising motifs I-IV shown in sticks and labeled (same view as Figure 2D). (E) Closeup of Shedu-ΔNL active sites in the octamer state. In the four protomers involved in octamer formation (left panel), the motif II loop is partially disordered and represented as a dotted line. The kinked ɑC helices for the two protomers making up the octamer interface (dark blue and yellow) are labeled, and the octamer interface itself is indicated with a dashed line. In the four protomers not involved in octamer formation (right panel), the active site is in an autoinhibited state equivalent to that observed in the full-length Shedu structure.

Compared to the full-length Shedu tetramer, the protomer-protamer packing across interface 1 in Shedu-ΔNL is looser, burying 1200 Å^2^ of surface area per protomer instead of the 1700 Å^2^ per protomer seen the full-length Shedu structure. This change, coupled with the kinking of the long αC helices in two protomers, enables the motif II loop in these two protomers to rearrange into a fully active state with D249 positioned for catalysis (**Figure 3D-E**). This rearrangement occurs only in the protomers that are involved in octamer formation, resulting in a total of four fully assembled active sites per Shedu-ΔNL octamer; in the four protomers (two per tetramer) not involved in octamer formation, these protomers’ active sites are in an autoinhibited state equivalent to that observed in the full-length Shedu structure (**Figure 3E**). The four active sites in an active conformation are arranged as two pairs, each roughly aligned across the octamer interface and positioned ∼40 Å apart from one another (**Figure 3C**). To test whether our structure of Shedu-ΔNL could represent an active state of Shedu, we purified Shedu-ΔNL after expression in an *in vitro* transcription/translation system and tested its activity on purified plasmid DNA. In contrast to the full-length *B. cereus* Shedu which showed negligible cleavage activity (not shown), Shedu-ΔNL showed robust DNA nicking and double-strand cleavage activity (**Figure 4A-B**).

**Figure 4.**
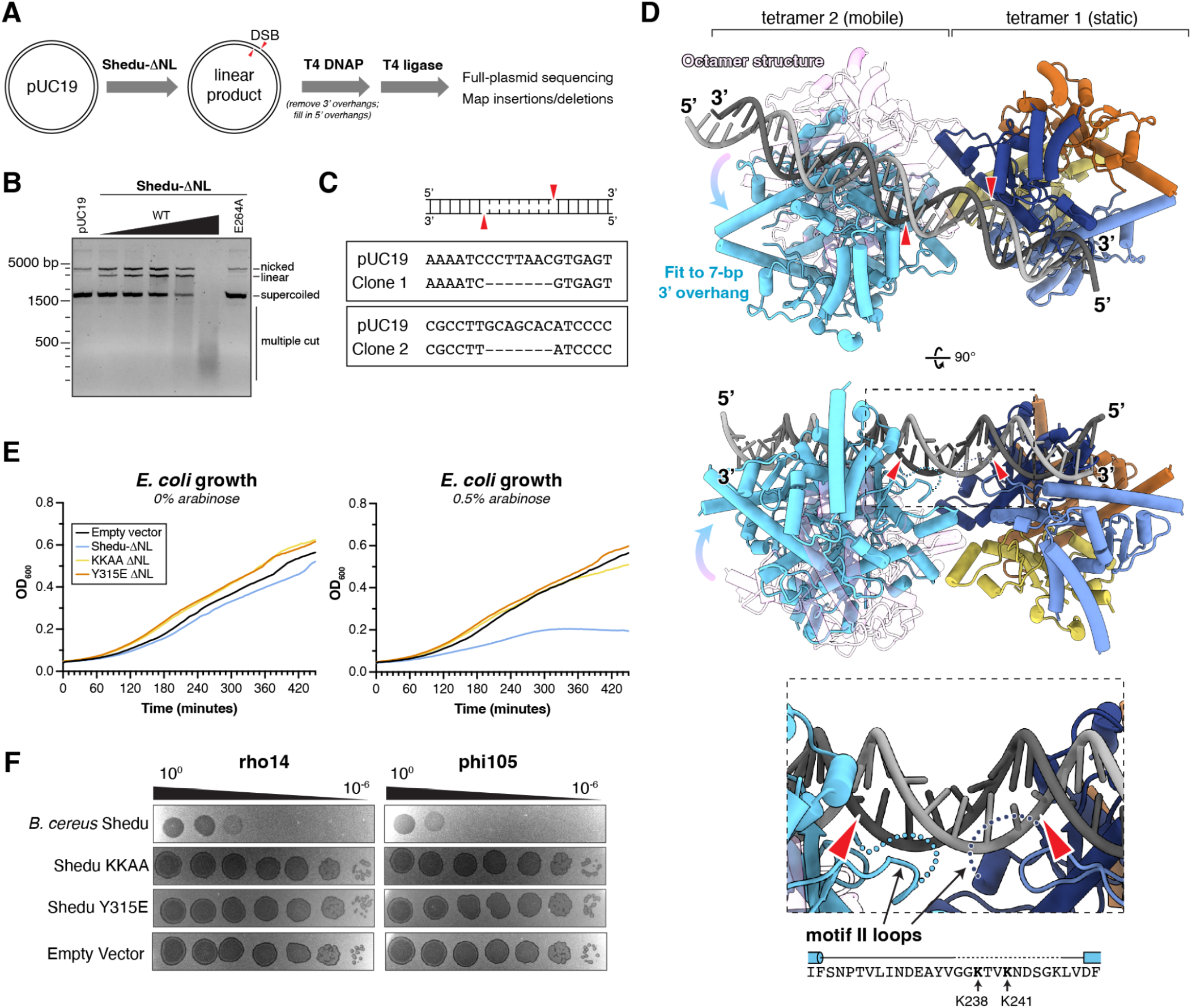
Shedu-ΔNL generates double-strand DNA breaks with 7-base 3′ overhangs. (A) Schematic of experiment to determine Shedu-ΔNL DNA cleavage geometry. “Blunting” by T4 DNA Polymerase (T4 DNAP) removes 3′ overhangs and fills in 5′ overhangs, which can be ligated by T4 DNA ligase, resulting in either deletions (for a 3′ overhang) or insertions (for a 5′ overhang) corresponding to the length of the respective overhang. DSB: double-strand DNA break. (B) Cleavage of pUC19 (2685 bp) by Shedu-ΔNL. The first lane has DNA alone; the next five lanes have added Shedu-ΔNL (3.125, 6.25, 12.5, 25, and 50 nM Shedu-ΔNL octamer); and the last lane has 50 nM Shedu-ΔNL E264A catalytic-dead mutant. (C) Full-plasmid sequencing results for two independent clones, after linearization by Shedu-ΔNL and blunting/ligation. Both show 7-bp deletions, indicating that the original DSBs generated 7-base 3′ overhangs. (D) Model of a DNA-bound Shedu-ΔNL octamer, generated by overlaying two Shedu-ΔNL tetramers on a structure of DNA-bound EndoMS (PDB ID 5GKE), then fitting each Shedu-DNA complex to an idealized double-stranded DNA (light gray/dark gray) such that two Shedu active sites are aligned to generate a 7-base 3′ overhang. Compared to the octamer conformation observed by cryoEM (light pink), the modeled conformation (cyan) requires a ∼40° rotation and ∼6 Å translation. This motion is likely accommodated by a combination of flexibility in the Shedu octamer interface and flexibility in the bound DNA. *Inset:* Closeup view of the proposed DNA cleavage sites (red arrowheads) and the location of the disordered Motif II loops, which contain two lysine residues (K238 and K241) mutated in the KKAA mutant. See **Supplemental Figure S5A** for sequence logo showing conservation of motif II loop residues. (E) Growth curves for *E. coli* transformed with arabinose-inducible vectors encoding *B. cereus* Shedu-ΔNL, in the absence of arabinose (left) or in the presence of 0.5% arabinose (right). Empty vector is shown in black, wild-type Shedu-ΔNL in blue, Shedu-ΔNL KKAA mutant (K238A/K241A) in yellow, and Shedu-ΔNL Y315E mutant in orange. See **Supplemental Figure S5B-E** for growth curves at other arabinose concentrations. Lines represent average values from triplicate measurements; see **Supplemental Figure S5B-E** for error bars. (F) Plaque assays with *B. subtilis* cells transformed with plasmids encoding *B. cereus* B4264 Shedu: wild-type, KKAA mutant, or Y315E mutant. Phages used were rho14 (left) and phi105 (right), with six ten-fold dilutions from a high-titer working stock. Images are representative of three independent trials. Images for wild-type Shedu and Empty Vector are the same as in Figure 1A.

While high concentrations of Shedu-ΔNL resulted in multiple nonspecific cleavage events as revealed by a smear on an agarose gel, we observed primarily nicks and single cleavage events at low protein concentration. To determine the specificity and geometry of DNA cleavage, we isolated linear plasmid DNA after Shedu-ΔNL cleavage, subjected the DNA ends to a blunting protocol using T4 DNA polymerase, then ligated and sequenced the resulting plasmids (**Figure 4C**). Out of seven sequenced plasmids, two showed a 7-bp deletion, consistent with Shedu-ΔNL generating a double-strand break with a 7-base 3′ overhang (**Figure 4C, Supplemental Table S2**). Five plasmids showed longer deletions ranging from 64 to 525 bp, suggestive of multiple cleavage events. There was no consistency in cleavage location among the seven sequenced plasmids, consistent with the enzyme’s DNA cleavage activity being sequence non-specific.

To determine how *B. cereus* Shedu might accomplish double-strand DNA cleavage and generate the observed 7-base 3′ overhang, we first generated a DNA-bound model of an individual active Shedu tetramer by overlaying one of the two activated protomers in a Shedu-ΔNL tetramer with DNA-bound EndoMS from *S. kodakarensis* ^28^. We next aligned two DNA-bound Shedu-ΔNL tetramer models on an ideal double-stranded DNA such that their active sites were positioned to cleave opposite strands and generate a 7-base 3′ overhang (**Figure 4D**). Strikingly, this model closely resembled that of the active Shedu-ΔNL octamer structure. A similar model of two Shedu tetramers aligned to generate a 6-base overhang resulted in significant clashes between the two tetramers, while a model of two Shedu tetramers aligned to generate an 8-base overhang showed the two tetramers rotated far apart from one another (not shown).

Rearrangement of the Shedu active site to position motif II residue D249 in the active site results in the C-terminal half of the motif II loop (residues 236-245) becoming disordered in our octamer structure (**Figure 4D**). We noticed that in our DNA-bound octamer model, this disordered loop is positioned such that it could participate in DNA binding. To test this idea, we generated a mutant with two lysine residues in the motif II loop mutated to alanine (K238A/K241A; KKAA). One of these lysine residues, K238, is part of a GGK motif that is highly conserved in close relatives of *B. cereus* Shedu (**Supplemental Figure S5A**). Consistent with a model in which these two lysine residues are important for DNA cleavage, the Shedu-ΔNL KKAA mutant was not toxic when expressed in *E. coli* (**Figure 4E, Supplemental Figure S5B-E**), nor was the full-length Shedu KKAA mutant able to protect against rho14 or phi105 infection (**Figure 4F**). Mutation of the key octamer interface residue tyrosine 315 to glutamate (Y315E) showed similar loss of toxicity in *E. coli* and loss of phage protection (**Figure 4E-F, Supplemental Figure S5B-E**). Altogether, our structure of Shedu-ΔNL and these functional data suggest that *B. cereus* Shedu is activated by octamer formation, which repositions motif II residue D249 and aligns pairs of active sites across the octamer interface for coordinated double-strand DNA cleavage. The fact that deletion of the protein’s N-terminal and linker domains resulted in a constitutively active enzyme, moreover, suggests that these domains inhibit Shedu activation until triggered by phage infection.

### Diversity of Shedu architecture suggests multiple activation mechanisms

To gain insight into Shedu regulation, we performed a deep sequence search for proteins containing the characteristic DUF4263 nuclease domain and identified 7,687 proteins, including the majority of the 1,246 examples identified by Doron et. al ^5^. Shedu homologs are mainly found in proteobacteria (53.3%), actinobacteria (16.7%), firmicutes (9.4%), bacteroidota (7.6%) and cyanobacteria (2.5%), plus a small number in archaea (36 instances in euryarchaeota, and 11 in DPANN archaea) (**Supplemental Table S3**). The proteins fall into two large groups based on their domain architectures: the majority (5,401 of 7,687), which we term Type 1 Shedu, possess one or more N-terminal domains in addition to the DUF4263 domain. Proteins in the smaller Type 2 Shedu group (2,286 of 7,687) possess only a DUF4263 domain, suggesting that these enzymes may be constitutively active (**Supplemental Table S3**).

We next investigated the N-terminal domains of Type 1 Shedu proteins in order to better understand their identity and complexity (**Figure 5A**). We first extracted the N-terminal domain sequences and used sequence-based clustering to generate 79 distinct clusters (**Supplemental Figure S6A**). 20 of these clusters were identifiable by sequence as known domain families in either the Pfam or CDD domain databases, leaving 59 unidentified domains that we termed N1 through N59. We used AlphaFold to predict the structures of all 79 N-terminal domains, then used structure-based clustering (**Supplemental Figure S6B**) and functional assignments from Pfam/CDD ^33,34^ or DALI searches ^35^ to group them into eight classes that we term Type 1A, 1B, 1C, 1D, 1E, 1F, 1G, and 1X (**Figure 5B, Supplemental Table S3**). Type 1A (1,684 instances) includes *B. cereus* Shedu and is characterized by sPC4/Whirly fold domains, all of which can be confidently modeled as homodimers (**Figure 5C**). As noted above, these domains may recognize single-stranded nucleic acids to sense phage infection. The next-largest two groups, Type 1B (1,594 instances) and Type 1C (307 instances), are characterized by novel ɑ/β domains with unknown functions, which we term NRF1 and NRF2 for N-terminal regulatory folds 1 and 2 (**Figure 5D-E**).

**Figure 5.**
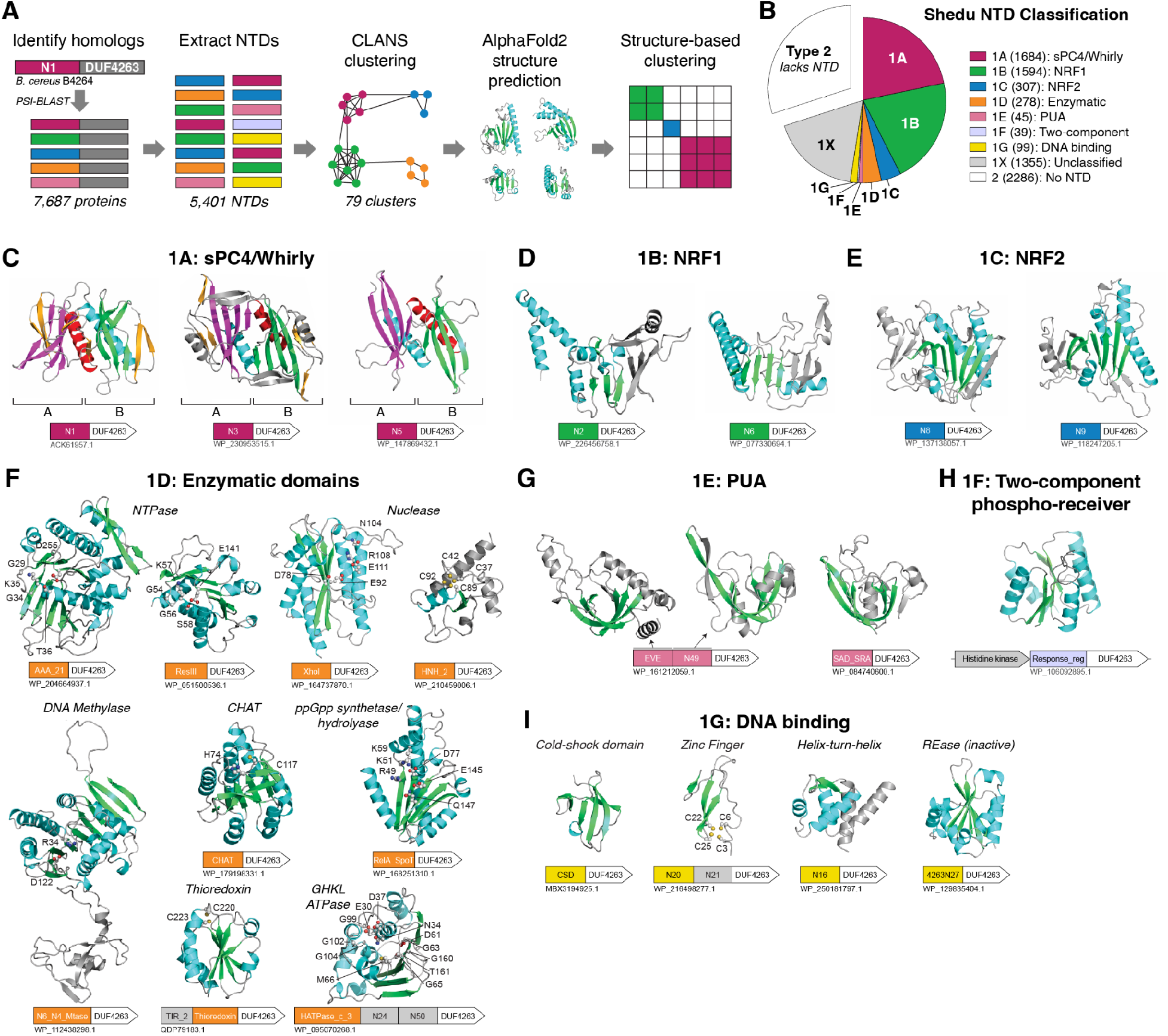
Shedu homologs encode diverse N-terminal regulatory domains. (A) Workflow for identification and classification of Shedu homologs’ N-terminal domains. See **Supplemental Figure S6A** for the CLANS analysis, and **Supplemental Figure S6B** for the DALI clustering analysis. (B) Pie chart showing the distribution of Shedu homologs into different classes. See **Supplemental Table S3** for full details. (C) Representative AlphaFold models and domain schematics of Type 1A Shedu homologs with NTDs related to sPC4/Whirly domains. Models show dimerized NTDs, with protomer A in red, purple, and yellow; and protomer B in cyan, green, and yellow. See **Supplemental Figure S7A** for additional examples. Each protein’s NCBI accession number is indicated. (D) Representative AlphaFold models and domain schematics of Type 1B Shedu homologs with NTDs of the NRF1 fold. See **Supplemental Figure S7B** for additional examples. (E) Representative AlphaFold models and domain schematics of Type 1C Shedu homologs with NTDs of the NRF2 fold. See **Supplemental Figure S7C** for additional examples. (F) Representative AlphaFold models and domain schematics of Type 1D Shedu homologs with NTDs predicted to have enzymatic activity. See **Supplemental Figure S7D** for additional examples. (G) Representative AlphaFold models and domain schematics of Type 1E Shedu homologs with NTDs of the PUA fold. Also see **Supplemental Figure S8A**. (H) Representative AlphaFold models and domain schematics of Type 1F Shedu homologs with NTDs related to two-component signaling systems’ phosphoreceiver domains. These proteins are often encoded adjacent to a predicted histidine kinase (see schematic). See **Supplemental Figure S8B** for additional examples. (I) Representative AlphaFold models and domain schematics of Type 1G Shedu homologs with NTDs predicted to bind DNA. See **Supplemental Figure S8C** for additional examples.

In addition to Type 1A, 1B, and 1C Shedu homologs, we identified a striking variety of different functional domains fused to the DUF4263 nuclease domain. We classified all Shedu homologs with putatively active enzymatic domains including NTPase, nuclease and methylase domains as Type 1D Shedu (278 instances; **Figure 5F, Supplemental Figure 7D**). Notable members of this class include enzymes with either DUF262 (PF03235) or N48, both of which are predicted NTPase domains belonging to the ParB/sulfiredoxin superfamily (**Supplemental Figure 7D**). The domain architectures of these Shedu homologs are similar to those of the known antiphage enzyme GmrSD/BrxU, also known as a Type IV R-M system, which contains a DUF262 domain and a DUF1524/HNH nuclease domain ^36,37^. DUF262 was recently demonstrated to regulate its associated nuclease domain through nucleotide binding and hydrolysis, in order to mediate antiphage immunity ^36^. Given the similarity of their domain architectures, we propose that Shedu proteins with NTPase NTDs may have a similar mechanism to GmrSD/BrxU. In other instances, the Shedu nuclease domain is associated with restriction endonuclease domains (ResIII, XhoI, Mrrcat) containing all requisite residues for nuclease activity (**Figure 5F, Supplemental Figure 7D**). Next, Shedu homologs with an active DNA methylase domain (N6_N4_MTase; PF01555) may be functional equivalent to Type II R-M systems, with the methylase domain being the modification component and the nuclease domain the restriction component. Finally, we identified Shedu homologs with several other enzymatic domains, including CHAT domains, putative ppGpp synthetase/hydrolase domains, thioredoxin folds, and GHKL ATPase folds (**Figure 5F, Supplemental Figure 7D**). Shedu homologs with a GHKL ATPase fold (HATPase_c_3) also generally possess a linked N24 domain, which is structurally related to the GHKL-associated middle domain of Hsp90-family chaperones, MORC proteins, and Type II DNA topoisomerase ATPase subunits (**Supplemental Table S3**).

Type 1E Shedu proteins (45 instances) possess domains belonging to the PUA (<u>P</u>seudo<u>U</u>ridine synthase and <u>A</u>rchaeosine transglycosylase) fold, including EVE, SAD_SRA and N49 (**Figure 5G, Supplemental Figure S8A**). Several PUA fold domains are known to recognize modified DNAs or RNAs ^38^; thus, these Shedus may be equivalent to novel type IV R-M systems that specifically target phages with modified genomes ^39^. Type 1F Shedu proteins (39 instances) possess putative two-component signaling system phosphoreceiver domains (Response_reg and N44), and are often encoded adjacent to a putative histidine kinase (**Figure 5H, Supplemental Figure S8B**). These proteins are likely activated by phosphorylation by their cognate histidine kinases, which in turn could respond to infection-related signals. Type 1G Shedu proteins (99 instances) encode various DNA binding domains at their N-termini, including winged helix-turn-helix, inactive restriction endonuclease, zinc finger, and cold-shock (CSD) domains (**Figure 5I, Supplemental Figure S8C**). Finally, Type 1X Shedu proteins (1,355 instances) encode domains that cannot be readily classified (**Supplemental Figure S8D**). Most of these domains are positioned at the N-terminus, but ∼12% (163 of 1,355) contain more than 100 amino acids C-terminal to the DUF4263 nuclease domain (**Supplemental Table S3**), suggesting that these proteins’s regulation may be distinct from the vast majority of Shedu proteins with N-terminal regulatory domains.

Overall, our sequence and structural analysis of Shedu proteins reveals a remarkably diverse set of putative regulatory domains that can likely sense and respond to a variety of infection- or stress-related signals. Our structures show that *B. cereus* Shedu – a Type 1A enzyme – is activated through the association of two tetramers into an octamer. We used AlphaFold 2 to predict the oligomeric state of Shedu proteins from all major families, including Type 2 Shedu proteins that lack an N-terminal domain, and found that in all cases these enzymes likely form homotetramers through their common DUF4263 nuclease domain (data not shown). In all cases, AlphaFold 2 also predicted an active site architecture matching our structure of the active *B. cereus* Shedu-ΔNL octamer, including the kinked ɑC helix, but could not model an octamer state of any Shedu protein (not shown). Thus, the question of which Shedu families may be activated by octamer formation remains open.

### Operonic organization of Shedu-containing loci

While most anti-phage immune systems encode separate infection sensor and effector proteins, Shedu is a single-gene immune system. We nonetheless identified several gene neighborhoods that were consistently associated with Shedu. Among 5,499 Shedu genes for which genomic information was available, we identified 602 examples of Shedu encoded within restriction-modification systems, including Type I, II, III, and IV systems (**Supplemental Figure S8E**). We also identified 87 examples of Shedu genes located within restriction-modification like BREX immune systems (**Supplemental Figure S8F**). These findings are consistent with prior work showing that Shedu is encoded in defense islands ^5^, and further suggest that Shedu may specifically complement the activity of restriction-modification or BREX systems. We also found that like many other defense island-associated immune systems, Shedu showed a strong association with transposable elements (1,110 instances). We further identified 121 instances of Shedu genes in prophage loci. The identification of Shedu genes in both transposable elements and prophages suggest that Shedu may enhance the fitness of these elements and their host cells by providing antiphage activities ^40,41^ and/or assisting superinfection exclusion.

## Discussion

The Shedu antiphage immune system is distinctive in that a single protein acts as both infection sensor and nuclease effector. Prior work ^5^ and our own data show that *B. cereus* B4264 Shedu strongly protects a *B. subtilis* host against two temperate phages, rho14 and phi105. We further find that the system functions non-abortively: that is, infected cells survive a phage infection rather than undergoing cell death to prevent the spread of infection. These results, and the fact that disrupting the protein’s predicted PD-(D/E)XK nuclease active site eliminates antiphage immunity, suggest that *B. cereus* Shedu directly recognizes these phages’ genomic DNA upon injection, and cleaves that DNA to eliminate the infection without harming the host cell’s genome (**Figure 6**).

**Figure 6.**
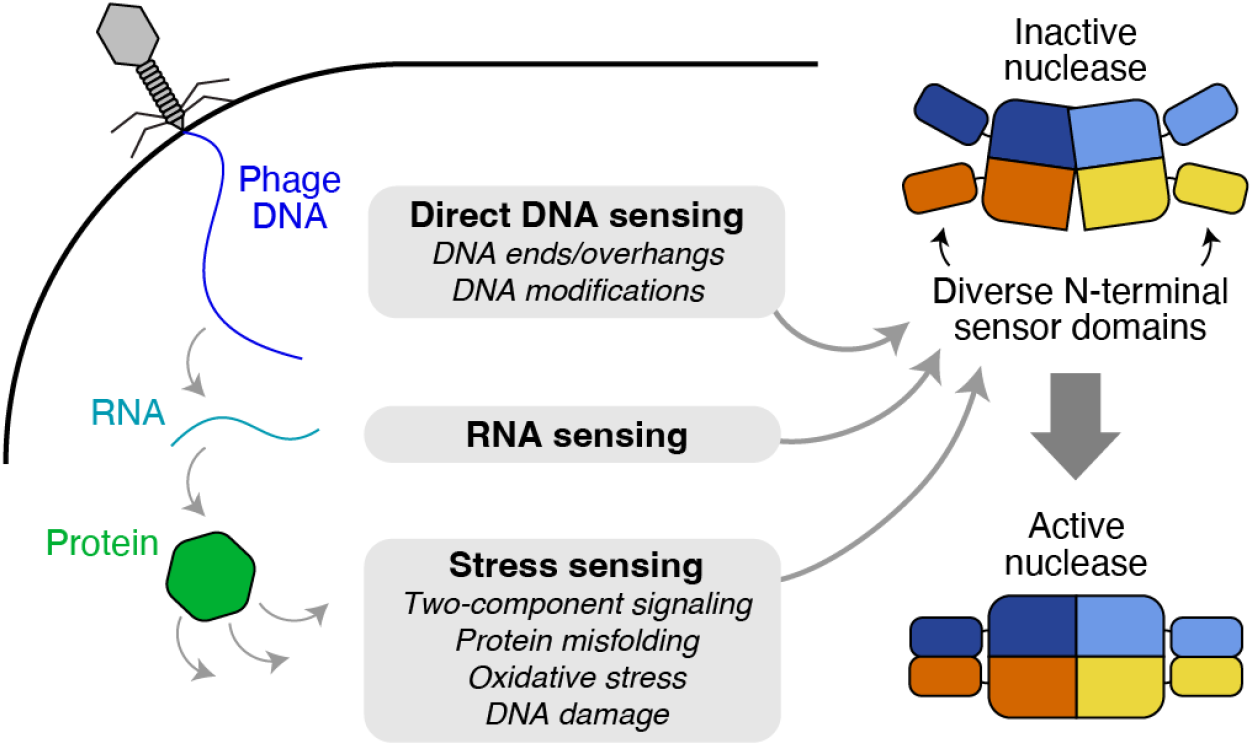
Model for Type 1 Shedu activation and immune function. Type 1 Shedu nucleases have diverse N-terminal domains that likely sense phage infection through a variety of signals. They may directly sense phage DNA by the presence of blunt or sticky DNA ends; they may sense phage RNA production, or they may sense a variety of stress signals including protein misfolding stress, oxidative stress, or DNA damage. The N-terminal domains may then directly recruit DNA substrates for cleavage (in direct DNA sensing enzymes) and/or allosterically activate the nuclease activity through changes in conformation and oligomerization state.

Our two cryoEM structures of *B. cereus* Shedu reveal the mechanistic basis for its regulation. We find that full-length Shedu assembles into a homotetramer through its C-terminal DUF4263 domain, which adopts a PD-(D/E)XK nuclease fold. In our structure of full-length Shedu, the nuclease active site is in an inactive conformation, due to the location of the extended motif II loop – which contains the critical motif II residue D249 – sequestered away from the active site. In addition, while the Shedu tetramer does share a common dimerization interface with canonical restriction endonucleases (interface 2 in Shedu), the active sites of the two subunits sharing this interface are not positioned to bind and cleave a double-stranded DNA. These findings explain our observation that full-length Shedu is non-toxic to cells even when overexpressed, and that it is inactive *in vitro*.

Deletion of the N-terminal and linker domains of *B. cereus* Shedu (Shedu-ΔNL) renders the enzyme highly toxic when expressed in *E. coli*. *In vitro*, the enzyme shows robust DNA nicking and double-strand DNA cleavage activity, and our data suggest that the two DNA strands are cut seven base pairs apart, leaving a 7-base 3′ overhang. Our cryoEM structure of Shedu-ΔNL reveals a homo-octamer assembled through association of two tetramers through an interface generated by kinking of the nuclease domains’ ɑC helices. In two subunits per tetramer, the extended ɑC helices kink by 90° to reveal a hydrophobic patch that forms a symmetric interaction with a second tetramer. Kinking of the ɑC helices also releases the motif II loop, which repositions residue D249 to complete the active site. Our modeling suggests that pairs of active sites arranged across the octamer interface can mediate coordinated cleavage of a double-stranded DNA to generate the observed 7-base 3′ overhang. Mutation of either Y315, a key residue in the octamer interface; or K238 and K241 in the motif II loop, which we propose aid DNA binding, eliminate toxicity of the Shedu-ΔNL protein and eliminate protection against phage infection. Thus, *B. cereus* Shedu is activated by formation of an octameric species, which is inhibited in the absence of infection by the protein’s N-terminal and linker domains.

The N-terminal domain of *B. cereus* Shedu is disordered in our cryoEM structure, but AlphaFold modeling suggests that these domains form dimers that extend from each end of the complex. The domain is predicted to adopt an sPC4/Whirly domain fold, which is known to interact with single-stranded nucleic acids ^30,31^. Mutation of a conserved arginine residue on the predicted nucleic acid binding surface of this domain eliminates antiphage immunity, indicating that this surface plays an important functional role. We propose that the N-terminal domain specifically recognizes phage-injected DNA, perhaps by binding the single-stranded end of the injected linear genome. Both rho14 and phi105 inject their DNA in a linear form with a 7-base 3′ overhang ^42,43^; the Shedu N-terminal domains may recognize this structure and then undergo a conformational change to enable Shedu octamerization and nuclease activation. One double-strand break near the end of the phage genome would eliminate the ability for the genome to circularize and replicate. Thus far, we have been unable to observe activation of full-length Shedu *in vitro* using DNAs mimicking this end structure (data not shown); thus the questions of what Shedu senses, and how sensing leads to nuclease activation, are important directions for future work.

Our initial sequence alignments of Shedu homologs revealed that these proteins’ C-terminal DUF4263 nuclease domains are well-conserved, but their N-termini are highly variable. Spurred by this observation, we undertook a comprehensive sequence- and structure-based analysis of Shedu homologs’ N-terminal domains across prokaryotes. We identified eight broad classes of Shedu proteins with a startling variety of both known and novel N-terminal domains. A minority of Shedu proteins (2,286 of 7,687) contain only a DUF4263 domain, and we term these Type 2 Shedu. These enzymes are likely constitutively active, and may recognize foreign DNA based on sequence or particular DNA modifications. Supporting this idea, a Type 2 Shedu from *Salmonella* was recently shown to protect an *E. coli* host against a phage T4 mutant unable to modify its genome, but this protection was dramatically reduced when the phage could modify its DNA to generate hydroxymethylcytosine or glucosyl-hydroxymethylcytosine ^27^. Growth curves of phage-infected cells showed that this protection was non-abortive, indicating that the phage DNA was likely targeted directly and that host DNA remained intact ^27^. Thus, Type 2 Shedu proteins may function similarly to restriction endonucleases, cleaving incoming phage DNA that is either unmodified or modified differently than that of the host.

The majority of Shedu proteins, which we term Type 1 Shedu, encode N-terminal domains that likely serve as infection or stress sensors and allosterically regulate their C-terminal DUF4263 nuclease domains (**Figure 6**). *B. cereus* Shedu falls into the largest class, Type 1A, which encodes sPC4/Whirly domains that likely recognize single-stranded nucleic acids. The next two largest classes, Type 1B and Type 1C, encode novel N-terminal domains that we term NRF1 and NRF2. These domains may recognize distinctive phage DNA structures or other infection-related signals. A Type 1B Shedu from *E. coli* (termed PD-T4-8; encodes an NTD in the N2 cluster) was recently shown to protect against phage T4, and was proposed to function by abortive infection ^7^. Other classes of Shedu proteins encode putative RNA binding (Type 1E) or DNA binding (Type 1G) domains, and/or enzymatic domains including DNA methylase, endonuclease, and NTPase domains (Type 1D). Some of these may bind particular DNA structures or modifications to sense infection. A class of GHKL ATPase-related NTDs, similar to Hsp90-family chaperones, may sense misfolded proteins, while thioredoxin-related NTDs may sense oxidative stress. Finally, a small class of Shedu proteins (Type 1F) encodes N-terminal phosphoreceiver domains typical of two-component signaling systems; these proteins are often encoded adjacent to a predicted histidine kinase. In canonical two-component systems, phosphorylation often induces dimerization of phosphoreceiver domains to mediate allosteric activation of fused enzymatic or signaling domains ^44^; a similar mechanism may apply to Shedu proteins with N-terminal phosphoreceiver domains.

Overall, our data reveal Shedu as a broad family of immune enzymes with a common nuclease core that is regulated by a remarkable variety of N-terminal sensory/regulatory domains. We have determined the mechanistic basis for activation of Type 1A Shedu, showing that its N-terminal domains allosterically regulate active site architecture and octamer formation. Future work will be needed to determine whether other families of Shedu proteins are activated in a similar manner, and to identify the diverse signals sensed by these enzymes’ N-terminal regulatory domains.

## Materials and Methods

### Cloning and Protein Purification

*Bacillus cereus* B4264 Shedu (NCBI accession ACK61957.1) full length gene was codon-optimized and synthesized (Integrated DNA Technologies), and cloned into UC Berkeley Macrolab vector 2-CT (Addgene #29706), which encodes an N-terminal TEV protease cleavable His_6_-MBP (maltose binding protein) tag. Mutant constructs were generated by PCR mutagenesis. Shedu-ΔNL constructs were cloned into UC Berkeley Macrolab vector 2-BT (Addgene #29666), which encodes an N-terminal TEV protease-cleavable His_6_ tag.

All proteins except wild type Shedu-ΔNL were recombinantly expressed in E. coli strain Rosetta2 pLysS (EMD Millipore) in 2x YT medium. Briefly, 1L cultures in 2L Erlenmeyer flasks were grown to an OD_600_ of 0.6 at 37°C, then the temperature was shifted to 20°C and IPTG (Isopropyl β-d-1-thiogalactopyranoside) was added to 0.2 mM to induce protein expression. Cultures were grown overnight (16-18 hours), then harvested by centrifugation.

For protein purification, two liters of cells were harvested, resuspended in resuspension buffer (25 mM Tris-HCl pH 8.5, 500 mM NaCl, 5 mM MgCl_2_, 10 mM imidazole, 5 mM β-mercaptoethanol, 5% glycerol), and lysed by sonication. The cell lysate was clarified with centrifugation for 30 min at 17,000 RPM, and the supernatant was loaded onto 2 ml Ni-NTA Superflow resin (Qiagen). The resin was washed once with 10 mL resuspension buffer and three times with 10 mL resuspension buffer supplemented with 30 mM imidazole, then eluted with 5 mL elution buffer (25 mM Tris-HCl pH 8.5, 300 mM NaCl, 5 mM MgCl_2_, 250 mM imidazole, 5 mM β-mercaptoethanol, 5% glycerol). 0.5 mg His_6_-tagged TEV protease ^45^ was added to the elution, and the mixture was dialyzed against dialysis buffer (10 mM Tris-HCl pH 8.5, 300 mM NaCl, 5 mM MgCl_2_, 5 mM β-mercaptoethanol, 5% glycerol) overnight at 4°C. TEV-cleaved proteins were run through a 5 ml HisTrap HP column (Cytiva), and flow-through fractions were combined with fractions from column washing with resuspension buffer supplemented with 40 mM imidazole. Combined fractions were concentrated to 500 µL and loaded onto a Superdex 200 Increase 10/300 column (Cytiva) equilibrated with size exclusion buffer (20 mM Tris-HCl pH 8.0, 300 mM NaCl, 5 mM MgCl_2_, 1 mM DTT, 5% glycerol). For structure determination, fractions were immediately used for cryoEM grid preparation (see below). For biochemical assays, fractions were concentrated, snap-frozen in liquid nitrogen, and stored at −80 °C.

Wild-type Shedu-ΔNL was toxic to E. coli and could not be expressed as above. For nuclease activity assays, wild type and E264A Shedu-ΔNL were expressed *in vitro* using an NEBExpress Cell-free *E. coli* Protein Synthesis System (New England Biolabs), then purified with Ni-NTA beads (Qiagen) using buffers described above.

### Oligomeric State Analysis by SEC-MALS

Protein oligomeric states were analyzed by SEC-MALS. 100 µl of purified proteins at ∼2 mg/ml concentration were injected onto a Superdex 200 Increase 10/300 column (Cytiva) on an HPLC (Agilent) connected to miniDAWN TREOS and Optilab T-rEX detectors (Wyatt Technology). The column was equilibrated overnight in size exclusion buffer. Molecular weight of different peaks were calculated using ASTRA version 6 or version 8 (Wyatt Technology).

### Negative Stain Electron Microscopy

For negative stain electron microscopy, purified full-length Shedu at 0.02 mg/mL in size exclusion buffer was applied to a glow-discharged carbon-coated copper grid, stained with three 1-minute incubations with filtered 2% uranyl formate, then blotted with filter paper and left to dry. Grids were imaged on an FEI Tecnai G2 Sphera microscope operating at 200 keV, fitted with a Gatan 2K x 2K CCD detector.

Micrographs were manually analyzed with ImageJ for particle measurements. For 2D class average calculations, 380 individual particles across 30 images were manually picked in cryoSPARC version 4^46^ and subjected to 2D class averaging. The class average displayed in **Figure 2C** was the most representative of individual particle images, and included 192 particles.

### Cryo Electron Microscopy

For grid preparation of full-length Shedu, freshly purified protein was collected from size exclusion chromatography and diluted to 18 µM (monomer concentration). Immediately prior to use, UltraAuFoil 1.2/1.3 300 mesh grids were plasma cleaned for 12 sec in a pre-set program using a Solarus II plasma cleaner (Gatan). 3.5 µl protein was applied to the grids in the environmental chamber of a Vitrobot Mark IV (Thermo Fisher Scientific) held at 4°C and 100% humidity. The grids were immediately blotted with filter paper for 5.5 sec with blot force setting of 20, followed by plunging into the liquid ethane cooled by liquid nitrogen. Grids were mounted into standard AutoGrids (Thermo Fisher Scientific) for imaging. The sample was imaged at the UCSD CryoEM facility using a Titan Krios G3 transmission electron microscope (Thermo Fisher Scientific) operated at 300 kV and configured for fringe-free illumination and equipped with a K2 direct electron detector (Gatan) mounted post Quantum 968 LS imaging filter (Gatan). The microscope was operated in EFTEM mode with a slit-width of 20 eV and using a 100 μm objective aperture. Automated data acquisition was performed using EPU (Thermo Fisher Scientific) and all images were collected using the K2 in counting mode. Ten-second movies were collected at a magnification of 165,000x and a pixel size of 0.65 Å, with a total dose of 54 e-/Å^2^ distributed uniformly over 40 frames. In total, 1311 movies were acquired with a realized defocus range of −0.8 to −2.0 μm.

Shedu-ΔNL grids were prepared similarly to full length protein, but were imaged at UCSD’s CryoEM facility using a Titan Krios G4 transmission electron microscope (Thermo Fisher Scientific) operated at 300 kV configured for fringe-free illumination and equipped with a Falcon IV direct electron detector with Selectris X energy filter. The microscope was operated in EFTEM mode with a slit-width of 20 eV and using a 200 μm objective aperture. Automated data acquisition was performed using EPU (Thermo Fisher Scientific) using EER format at a magnification of 130,000x and a pixel size of 0.935 Å, with a total dose of 55 e-/Å^2^ and rate of 5 eps. In total, 1825 movies in flat, and 1708 movies with 30° tilt were collected with a realized defocus range of −0.8 to −2.2 µm.

CryoEM data analyses were performed in cryoSPARC version 3.2 ^46^. Movies were motion-corrected using patch motion correction (multi) and CTF-estimated using patch CTF estimation (multi). For the full length Shedu sample, initial particles were picked with blob picking in cryoSPARC with diameter of 58 to 160 Å, and over 2 million particles were picked. A small subset of particles were subjected to two-dimensional classification, and revealed classes that look like Shedu protein complexes. Those classes were used as templates for template picking, which resulted in 954,170 particles. Several two-dimensional classifications and heterogeneous refinements were performed to clean up the particles, then *ab initio* structure determination was performed. Finally, 149,464 particles were sorted and refined using the non-uniform refinement method in cryoSPARC ^47^ with the following options enabled: C1 symmetry; minimize over-particle scale; optimize per-particle defocus; optimize per-group CTF params. This resulted in a tetramer Shedu complex reconstruction with a resolution of 2.9 Å using the 0.143 Fourier Shell Correlation cutoff between masked independently refined half-maps. Extensive trials with larger box and particle sizes were attempted in order to resolve the full ∼21 nm long particle including N-terminal and linker domains, without success.

For Shedu-ΔNL (E264A), untilted and tilted data were processed in parallel. Circular blob picking with diameter 50 to 100 Å was performed to identify over 2 million particles in total. Over a million particles from the untilted dataset went through several two-dimensional classification and heterogeneous refinement to get a final 276,404 particles (with 4x binned), which was refined to a coarse structure using homogeneous refinement. Two-dimensional templates were generated and 10 of them were used as templates to run template-based particle picking. The newly template-picked particles were cleaned and classified similarly to get a new particle set of 167,946 particles. Around a million particles from blob picking in the tilted data were cleaned and classified similar to the untilted dataset, which resulted in 170,539 particles. At this stage, particles of both datasets were combined and used to refine an octamer structure using homogeneous refinement to align all particles. Due to the flexibility of the octamer interface, we performed local refinement on one tetramer, with fulcrum location set at box center and non-uniform refinement enabled. This led to a 2.77 Å resolution map of one tetramer within the octamer. For the octamer structure refinement, we took the combined 331,076 particles, further cleaned with two-dimensional classification, which resulted in 260,388 particles. These particles were aligned with homogeneous refinement, and then 3D variability analysis was performed with 3 models and with filter resolution set at 4.5 Å. One model showed continuous movement of structure. This model was resolved with a 3D display method in an intermediate mode, with 11 frames. The first 4 frames of particles (138,326 particles) were refined with non-uniform refinement with the following options enabled: C1 symmetry, optimize per-particle defocus; optimize per-group CTF params.

An initial model of full length Shedu tetramer was generated using AlphaFold2 ^48,49^. This model was manually docked into the Shedu full length final cryo-EM map using UCSF Chimera ^50^ and rebuilt in COOT ^51^. For Shedu-ΔNL, the initial model was built against a locally refined map of the tetramer using the AlphaFold2 model. Then an octamer was built in COOT by expansion of the tetramer model. Both the Shedu full length tetramer and the Shedu-ΔNL octamer models were real-space refined in phenix.refine ^52^. Structure validation was performed with MoProbity ^53^ and EMRinger ^54^. Structures were visualized in ChimeraX ^55^ and PyMOL (Schrödinger).

### *B. subtilis* strain construction

*B. subtilis* BEST7003 cells were used as a host. The native genomic locus of *Bacillus cereus* B4264 Shedu (NCBI accession NC_011725.1, bases 954720-956931) ^5^ was synthesized (GeneArt/Invitrogen) and cloned into pSG1 vector between AscI and NotI sites. Mutants were generated by PCR mutagenesis. For transformation, 10 μl of a saturated overnight culture of *B. subtilis* were diluted into 1 ml of MC medium (80 mM K_2_HPO_4_, 30 mM KH_2_PO_4_, 2% glucose, 30 mM trisodium citrate, 22 μg/mL ferric ammonium citrate, 0.1% casein hydrolysate (CAA), 0.2% potassium glutamate) supplemented with 10 μl 1M MgSO_4_. After 3 hours of incubation (37°C, 200 RPM), 300 μl was transferred to a new 15 ml tube and ∼200 ng of plasmid DNA was added, and then incubated for another 3 hours. 300 µlL of transformation culture was plated on an LB agar plate supplemented with spectinomycin. Successful transformation would integrate the Shedu locus along with spectinomycin resistance gene into the *amyE* locus. Colonies were PCR screened to verify the integration of the Shedu locus on the *B. subtilis* genome.

### Phage plaque assays

Phages phi105 and rho14 were ordered from Bacillus Genetic Stock Center (#1L11 and 1L15). Phages were recovered from filter paper by soaking on a *B. subtilis* lawn overnight, and single plaques were isolated from re-streaking onto new *B. subtilis* lawns. Phage genomes from stocks isolated from single plaques were verified with full-genome sequencing prior to use (SeqCenter, LLC). Phage amplification was performed with freshly grown *B. subtilis* liquid culture in LB media, grown to OD_600_=0.4. *B. subtilis* cells were diluted to OD_600_=0.1 in LB media supplemented with 10 mM MgCl_2_ and 2 mM CaCl_2_, then infected with 1-to-100 ratio of stock phage for 4 hours at 30°C. Amplified phages were harvested through spinning down the cells/cell debris and filtered with 0.2 µM filter. Amplified phage titres (PFU/ml) were determined by plaque counting on a B. subtilis lawn on an LB top agar plate (LB media + 0.35% agar, supplemented with 10 mM MgCl_2_ and 2 mM CaCl_2_).

For phage plaque assays, we used a modified double-layer agar plate technique ^56^. Overnight cultures of *B. subtilis* were diluted into 5 ml fresh media and incubated until the OD_600_ reached 0.2. 500 µl of cells were mixed with 4.5 ml top agar media (LB media + 0.35% agar) supplemented with 10 mM MgCl_2_, 2 mM CaCl_2_ and 50 µg/ml spectinomycin, and poured on top of an LB agar plate containing spectinomycin. The plate was incubated 45 minutes to allow the top agar to solidify and slightly dry before phage spotting. Each phage stock is approximately 10^10 PFU/mL, and stocks were serially diluted (10-fold dilutions) starting from a 10-fold dilution of the original stock, then 3 µl of each dilution was spotted on each plate. Plates were incubated at 30°C overnight before scoring plaques.

We performed plaque assays with phage Phi29, a third phage that *B. cereus* Shedu was reported to show protection against ^5^. We were, however, unable to detect any protective effect of Shedu against this phage (not shown).

### Nuclease Assays

Nuclease assays were performed with pUC19 plasmid (2685 bp). B. cereus Shedu-ΔNL at the indicated concentrations was mixed with 150 ng plasmid in reaction buffer (20 mM HEPES pH 7.0, 100 mM NaCl, 5 mM MgCl_2_, and 1 mM DTT) for 2 hours. Digested samples were analyzed by electrophoresis on 0.5x TBE agarose gel. Gels were stained with ethidium bromide and imaged by UV illumination.

To determine double-strand DNA cleavage geometry, linearized pUC19 from incubation with Shedu-ΔNL was gel-extracted and purified from a well-resolved TBE agarose gel. T4 DNA polymerase (NEB) treatment in the presence of 500 µM dNTPs was performed at 12°C for 15 min, then quenched with 10 mM EDTA and incubation at 75 °C for 20 min. The reaction mixture was further purified with a PCR clean up kit (Qiagen). 0.5 µL T4 DNA ligase (New England Biolabs) was added into a 10 µl reaction mixture, and the ligation reaction was performed at room temperature for 2 hours. Half of the reaction mixture was used to transform Novablue cells (EMD Millipore Sigma), and the amplified plasmids from 7 colonies were mini-prepped and subjected to full-plasmid sequencing (Primordium Labs).

### Toxicity Assays

Toxicity assessment of Shedu (full length and ΔNL) was performed with an arabinose-inducible expression system in *E. coli*. Briefly, genes were cloned into a modified pBAD vector (UCB Macrolab vector 8B, Addgene #37502). Mutants were made by PCR mutagenesis, and all constructs were verified by sequencing. Plasmids containing various Shedu constructs were transformed into Novablue cells (EMD Millipore Sigma) and grown overnight on an LB plate supplemented with 50 µg/ml carbenicillin. A single colony was picked and grown in LB plus carbenicillin overnight, then diluted to OD_600_=0.1 in fresh media in the morning. After growing at 37°C to an OD_600_ of 0.5, they were diluted again to OD_600_=0.05 in LB media supplemented with 50 µg/ml carbenicillin and with indicated amounts of arabinose. 100 µl were added to each well of a standard clear 96-well plate with lid (Corning). The plate was incubated in a plate reader (Tecan Sunrise) at 30°C and OD_600_ readings were taken every 5 minutes for 100 cycles in total, with medium shaking for 200 seconds in between. Each condition was performed as a triplicate on a single plate.

For growth curves with phage infection, *Bacillus subtilis* cells integrated with wild type Shedu or mutants were grown in liquid culture overnight from a glycerol stock. The next morning, the cultures were inoculated into fresh LB media supplemented with spectinomycin. When the cultures reached an OD_600_ of 0.5, cells were diluted to OD_600_=0.1 in fresh media. 100 µl of diluted cultures were added to each well of a standard clear 96-well plate with lid (Corning), then 5 µL of phages at different dilutions were added to achieve the indicated multiplicity of infection (MOI) values. Plates were incubated in a plate reader (Tecan Sunrise) at 30°C and OD_600_ readings were taken every 5 minutes for 100 cycles in total, with medium shaking for 200 seconds in between. Each condition was performed as a triplicate on a single plate.

### Bioinformatics

To obtain a comprehensive collection of Shedu homologs, we used the DUF4263 domain of *Bacillus cereus* B4264 Shedu (NCBI accession ACK61957.1) as the query in PSI-BLAST (Position-Specific Iterated BLAST), searching against the NCBI NR database ^57^. The search was performed iteratively with a cut-off e-value of 0.001 until convergence was reached, which resulted in the retrieval of 7,687 protein homologs containing the DUF4263 domain. A custom Python script was used to extract the sequences N-terminal to the DUF4263 domain of each sequence, with N-terminal domain sequences shorter than 40 amino acids discarded. The CD-HIT program ^58^ was utilized to remove highly similar sequences (c=0.97), which led to a collection of 3941 unique N-terminal domain sequences. These sequences were subjected to the CLANS analysis ^59^, which uses a Fruchterman and Reingold graph drawing algorithm to cluster the sequences based on the presence of significant high-scoring segment pairs (HSPs) detected by all-against-all BLASTP searches (P-value>0.0001). For any cluster of sequences, segregated as a dense subgraph, one or two representative sequences were selected for structural modeling using AlphaFold 2 ^48,49^ and domain dissection was performed based on sequences’ inter-residue distance matrices. Domain annotation was performed using HMMSCAN ^60^ searching against PFAM profiles ^34^ and our own custom domain profiles. For previously unrecognized domain families, homologous sequences were collected using PSI-BLAST searches, clustered by BLASTCLUST, a BLAST score-based single-linkage clustering method(ftp://ftp.ncbi.nih.gov/blast/documents/blastclust.html), to remove highly similar sequences, and multiple sequence alignments (MSAs) were built using the PROMALS3D program ^61^.

To classify N-terminal domains, representative AlphaFold2 models were selected and combined for all-against-all structural similarity comparisons using DALI ^62^. The structures were further clustered via average linkage hierarchical clustering based on a DALI Z-score matrix ^63^. In this analysis, we used the dimeric structure of novel Spc4/Whirly domains modeled by AlphaFold2 for DALI comparisons as DALI cannot recognize the similarity between the two monomer types of the Spc4/Whirly domains.

Additionally, DALI structure similarity searches against the PDB database were conducted to reveal distant relationships of novel domains and any other known structures. Electrostatic potential of protein surfaces was calculated using the Adaptive Poisson-Boltzmann Solver (APBS) software ^64^. Visualization and analysis of protein structures was conducted using PyMOL (http://www.pymol.org) or ChimeraX ^55^.

Gene neighborhood analysis ^65^ was conducted for all the 7687 Shedu proteins by collecting their upstream and downstream genes (seven genes on each side for the majority of cases; more genes retrieved for cases with large operons). Then, both the Shedu proteins and proteins encoded by neighbor genes were clustered based on sequence similarities via BLASTCLUST. Each cluster and its respective proteins were then annotated by their domain architectures via the HMMSCAN program described above.

## Acknowledgements

The authors thank Rotem Sorek for providing *B. subtilis* cells, pSG1 vector, and experimental advice; and members of the Corbett and Zhang labs for helpful conversations and critical reading of the manuscript. K.D.C. acknowledges support from the National Institutes of Health (R35 GM144121). D.Z. acknowledges support from Saint Louis University. The authors acknowledge the facilities, along with the scientific and technical assistance of the staff of the CryoEM facility at UC San Diego.

## Declaration of Interests

The authors declare no competing interests.

---

**Figure S1.**
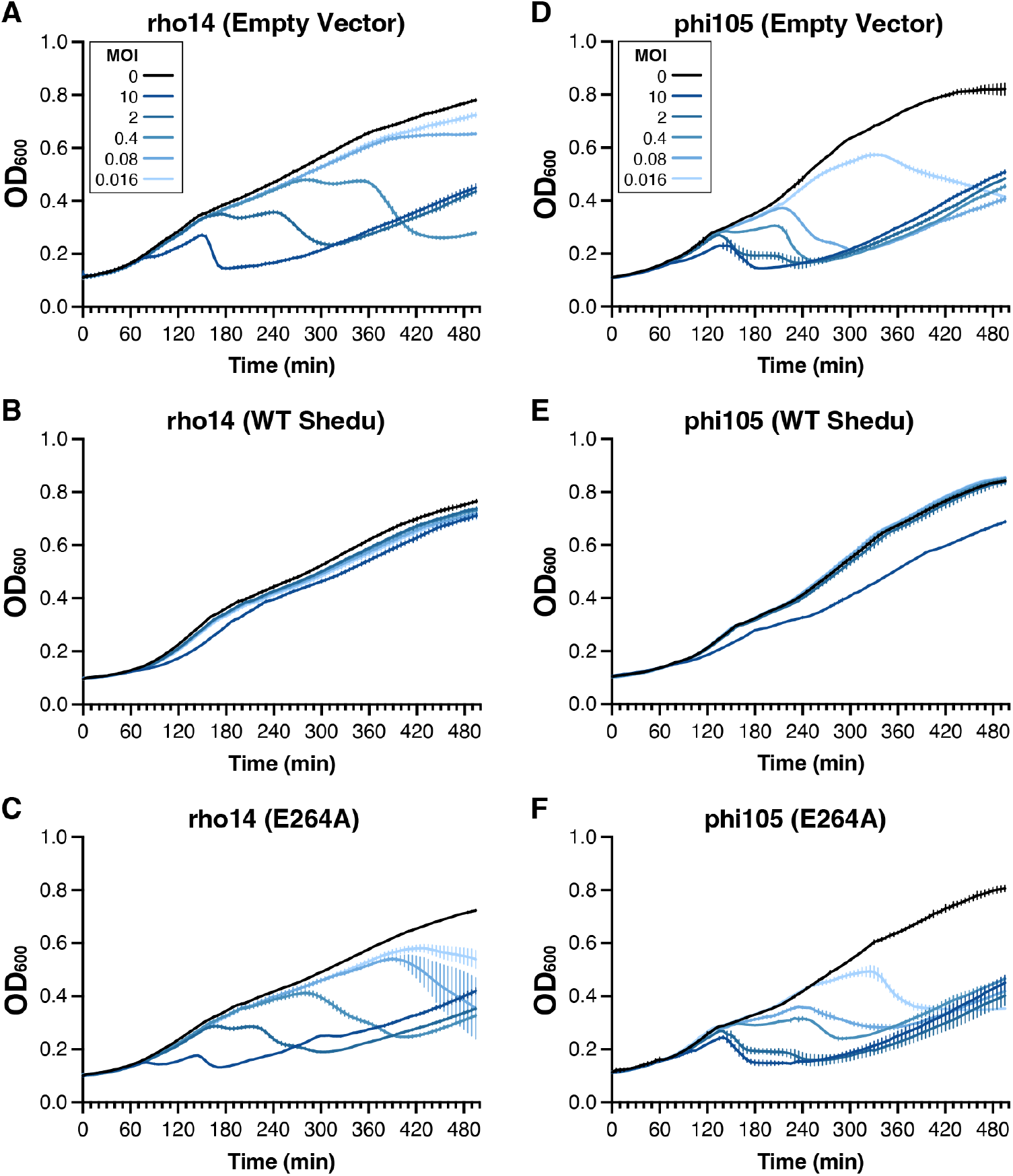
*B. cereus* Shedu is a non-abortive immune system. Growth curves of *B. subtilis* cells transformed with plasmids encoding *B. cereus* B4264 Shedu (empty vector: panels A and D; wild-type: panels B and E: E264A: panels C and F) with rho14 (panels A-C) or phi105 (panels D-F) at different multiplicity of infection (MOI) values from 10 to 0.016 (five-fold dilutions). Lines and error bars represent average values and standard deviation from triplicate measurements.

**Figure S2.**
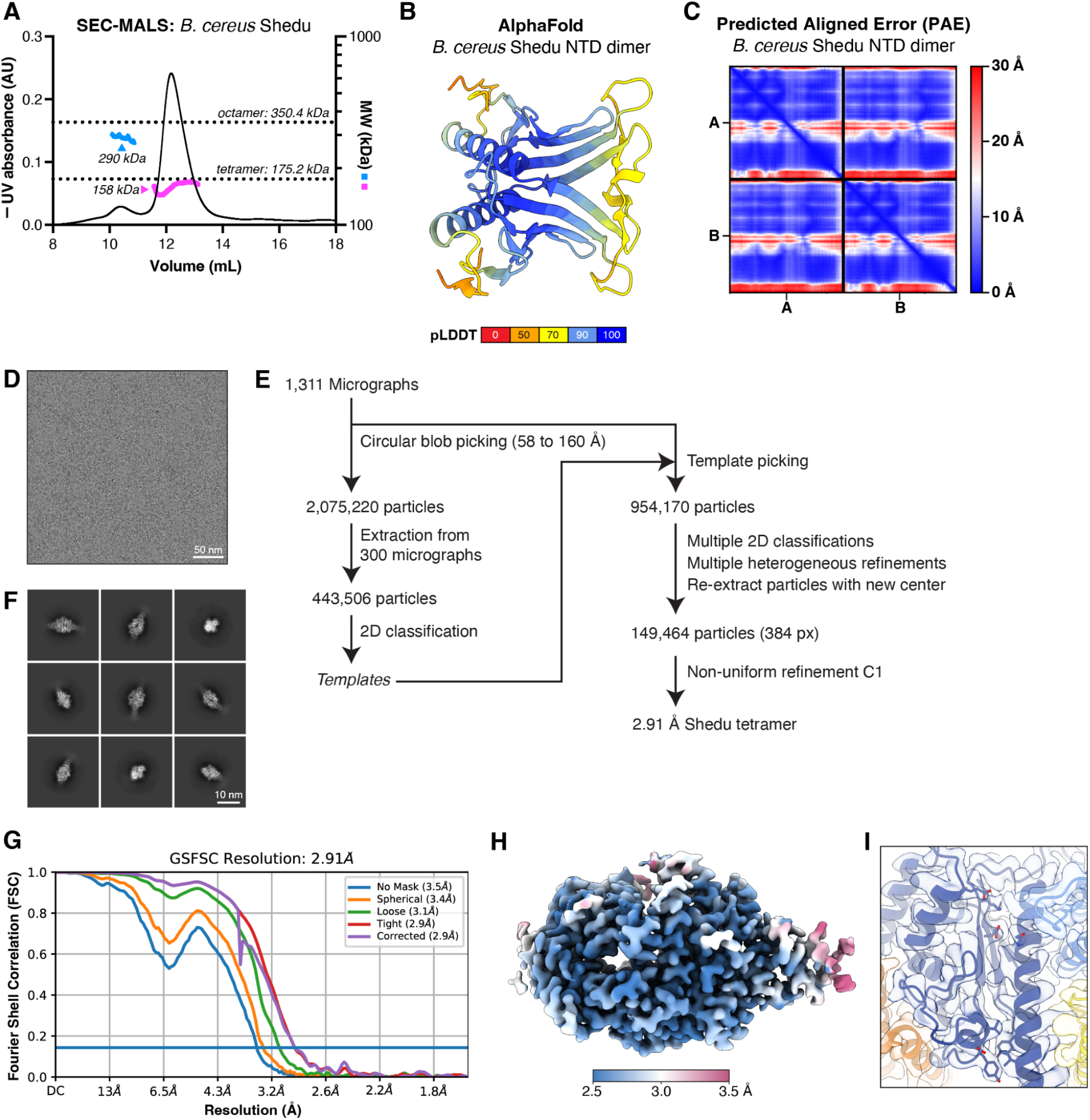
Cryoelectron Microscopy workflow for FL tetramer. (A) SEC-MALS analysis of full-length *B. cereus* Shedu. Measured molecular weights (blue and pink) are indicated, and the theoretical molecular weight of tetramer and octamer species are indicated with dotted lines. (B) AlphaFold 2 predicted structure of a *B. cereus* Shedu NTD dimer, colored by pLDDT confidence score. (C) Predicted Aligned Error (PAE) plot for AlphaFold 2 structure prediction of a *B. cereus* Shedu NTD dimer. (D) Representative cryoEM micrograph of full-length *B. cereus* Shedu. (E) Workflow for cryoEM structure determination of full-length *B. cereus* Shedu. (F) Representative 2D classes. (G) Fourier Shell Correlation (FSC) curves for the final refinement of full-length *B. cereus* Shedu. (H) Final cryoEM density, colored by local resolution from 2.5 Å (blue) to 3.5 Å (pink). (I) Sample cryoEM density (semitransparent surface) for Shedu surrounding the nuclease active site. View equivalent to Figure 2D.

**Figure S3.**
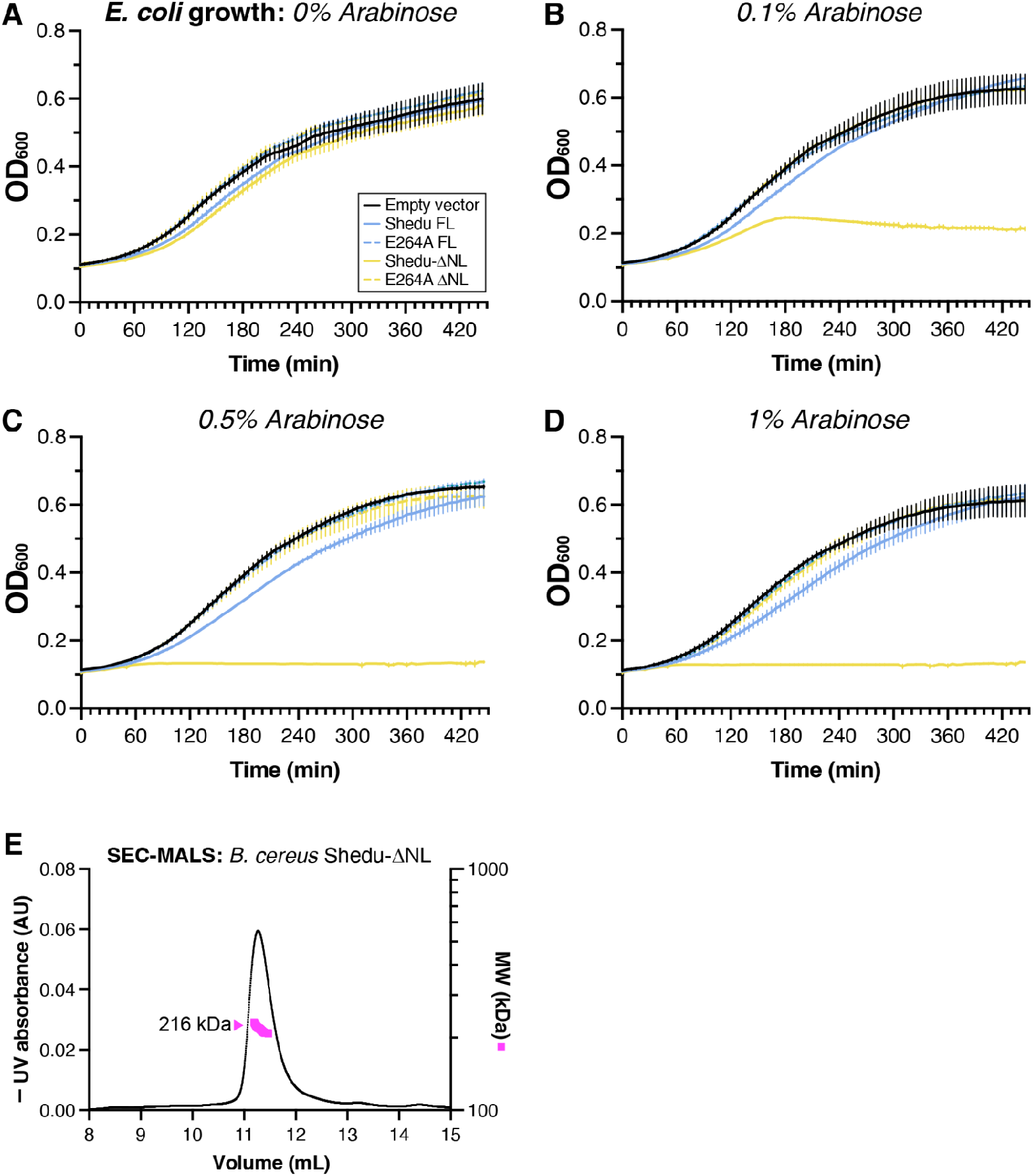
Shedu N-terminal domains regulate nuclease activity. (A) through (D) Growth curves for *E. coli* transformed with arabinose-inducible vectors encoding *B. cereus* Shedu, in the absence of arabinose (A) or in the presence of 0.1 (B), 0.5% (C), or 1% arabinose (D). Empty vector is shown in black, full-length Shedu in blue, and Shedu-ΔNL in yellow. E264A mutants for full-length Shedu and Shedu-ΔNL are shown as dashed lines. Lines and error bars represent average values and standard deviation from triplicate measurements. (E) SEC-MALS analysis of *B. cereus* Shedu-ΔNL. Measured molecular weights (pink) is indicated, the theoretical molecular weight of an octamer species is 210 kDa.

**Figure S4.**
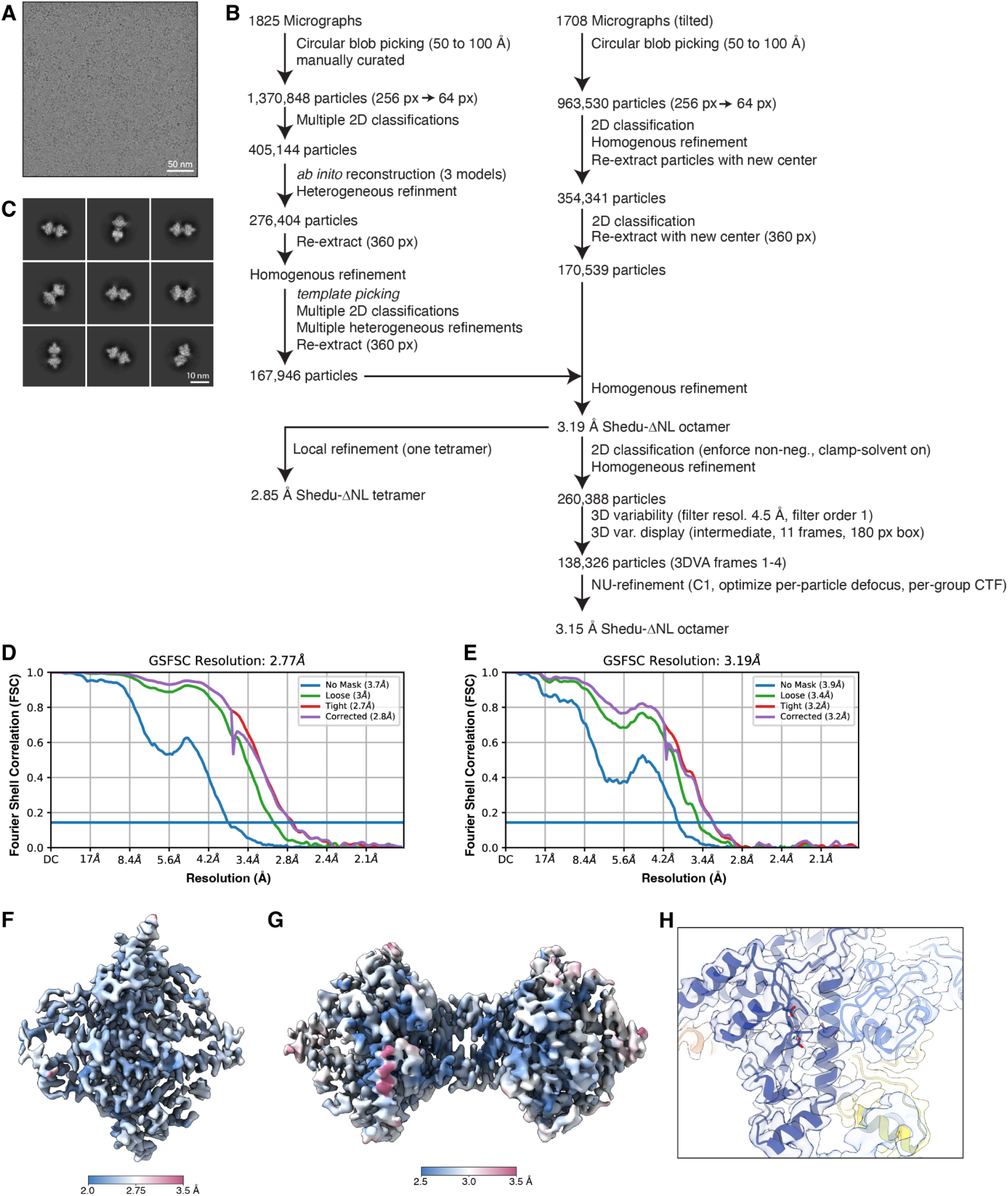
Cryoelectron Microscopy workflow for Shedu-ΔNL octamer. (A) Representative cryoEM micrograph of *B. cereus* Shedu-ΔNL. (B) Workflow for cryoEM structure determination of Shedu-ΔNL. (C) Representative 2D classes. (D) Fourier Shell Correlation (FSC) curves for the final refinement of one tetramer within the Shedu-ΔNL octamer. (E) Fourier Shell Correlation (FSC) curves for the final refinement of the full Shedu-ΔNL octamer. (F) Final cryoEM density for the final refinement of one tetramer within the Shedu-ΔNL octamer, colored by local resolution from 2.0 Å (blue) to 3.5 Å (pink). (G) Final cryoEM density for the final refinement of the full Shedu-ΔNL octamer, colored by local resolution from 2.5 Å (blue) to 3.5 Å (pink). (H) Sample cryoEM density (semitransparent surface) for Shedu-ΔNL surrounding the nuclease active site. View equivalent to Figure 3D.

**Figure S5.**
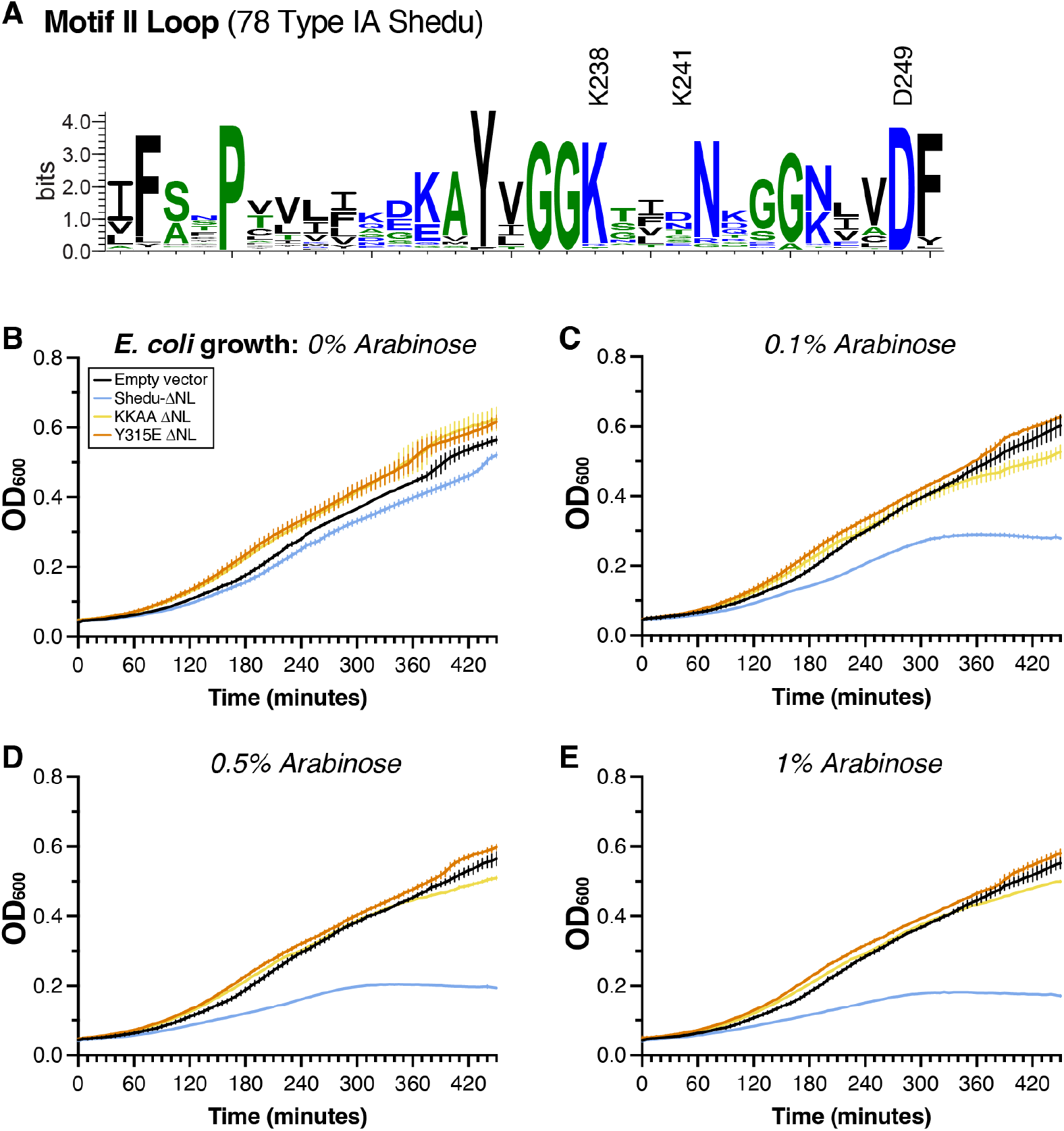
The *B. cereus* Shedu octamer is the active species. (A) Sequence logo for the Shedu motif II loop, created from a sequence alignment of 78 Type I Shedu proteins related to *B. cereus* Shedu. The two lysine residues mutated in the KKAA mutant (K238 and K241) are labeled, as is motif II residue D249. (B) through (E) Growth curves for *E. coli* transformed with arabinose-inducible vectors encoding *B. cereus* Shedu-ΔNL, in the absence of arabinose (B) or in the presence of 0.1 (C), 0.5% (D), or 1% arabinose (E). Empty vector is shown in black, wild-type Shedu-ΔNL in blue, KKAA mutant (K238A/K241A) in yellow, and Y315E mutant in orange. Lines and error bars represent average values and standard deviation from triplicate measurements.

**Figure S6.**
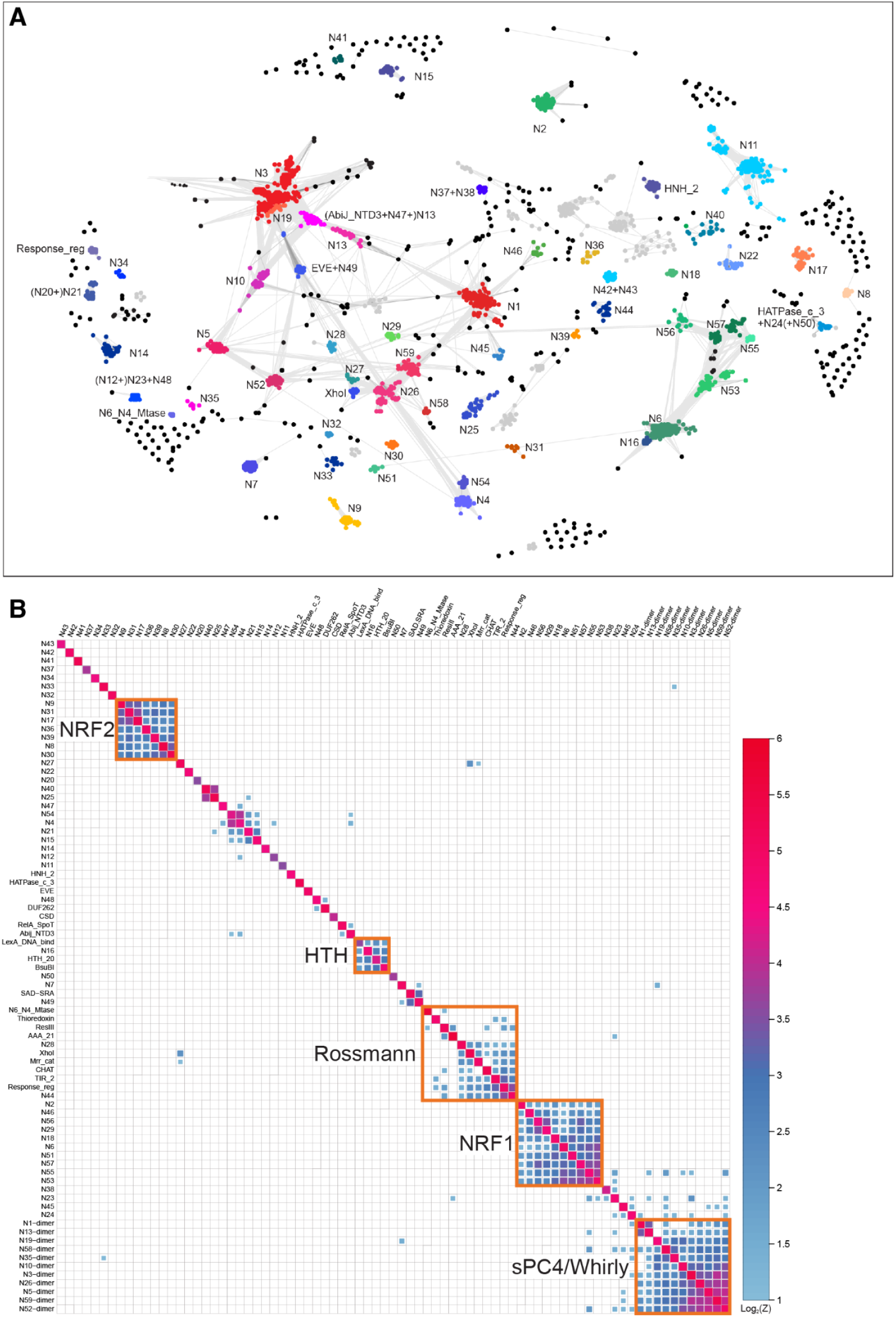
Shedu homologs encode diverse N-terminal regulatory domains (part 1) (A) CLANS sequence-based clustering of isolated Shedu N-terminal domains. Identified clusters are colored and labeled. See **Supplemental Table S3** for full list and classification. (B) DALI-based structural clustering of AlphaFold models of each Shedu NTD. Clusters representing common folds are outlined in orange and labeled.

**Figure S7.**
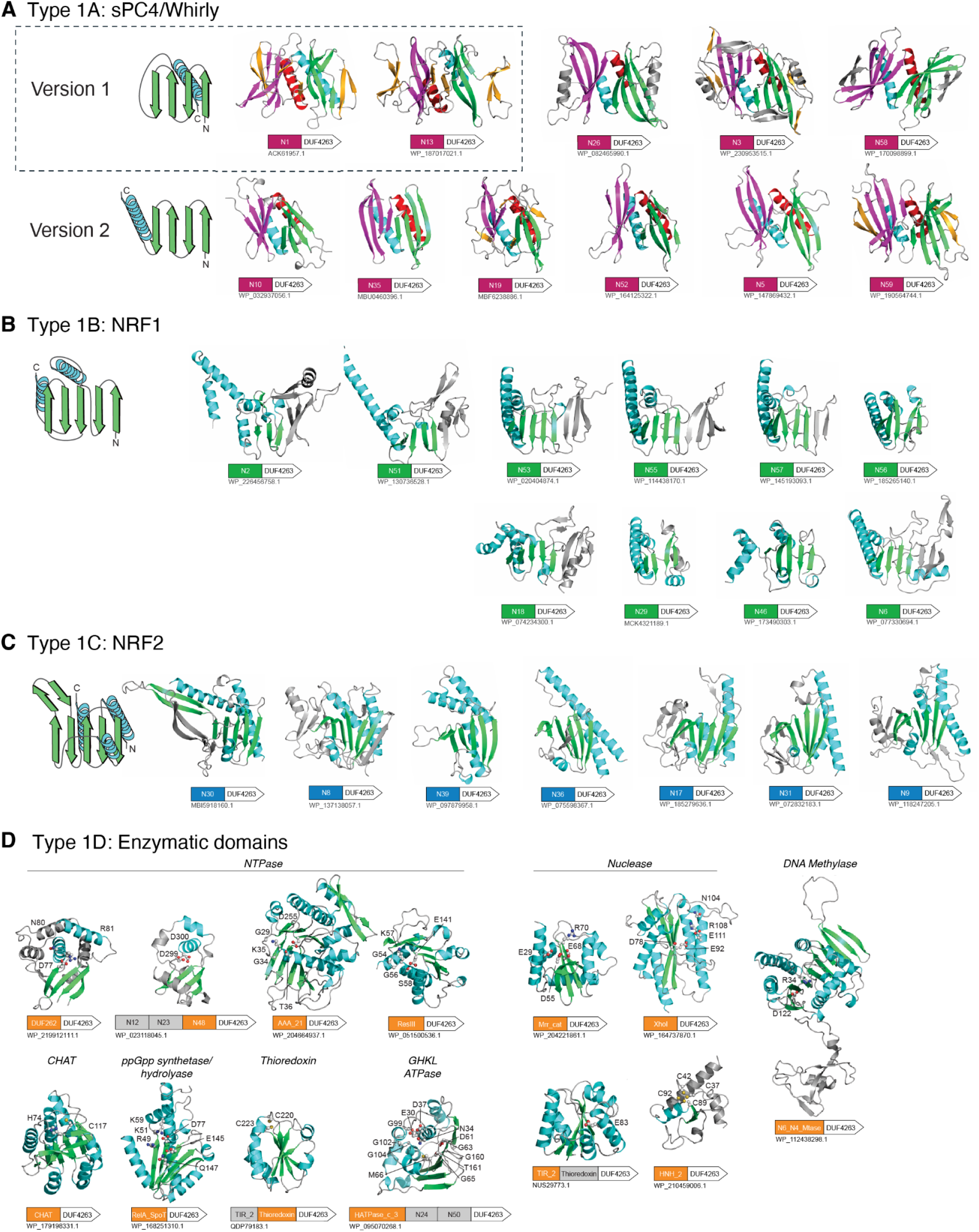
Shedu homologs encode diverse N-terminal regulatory domains (part 2) (A) Representative AlphaFold models and domain schematics of Type 1A Shedu homologs with NTDs related to sPC4/Whirly domains. All members of this group can be confidently modeled as homodimers by AlphaFold; models are colored with protomer A in red, purple, and yellow; and protomer B in cyan, green, and yellow. In version 1 of this fold (dotted box), the C-terminal α-helix of each protomer is packed against its own β-sheet. In version 2, the α-helix of each protomer is packed against the β sheet from the opposite protomer. Each protein’s NCBI accession number is indicated. (B) Representative AlphaFold models and domain schematics of Type 1B Shedu homologs with NTDs of the N-terminal Regulatory Fold 1 (NRF1) family. These domains share a previously undefined structural core, consisting of β_1_β_2_α_1_β_3_β_4_β_5_α_2_. No structural homologs can be retrieved in PDB by the DALI server, but they show significant structural similarity to one another in all-against-all comparisons (**Figure S6B**). For all examples except N46, a highly positively charged cavity can be formed by the two core α-helices, suggesting that these domains may also bind to nucleic acids. (C) Representative AlphaFold models and domain schematics of Type 1C Shedu homologs with NTDs of the NRF2 family. This previously unidentified fold has three layers with a central five-stranded β sheet connected by three α-helices on the two sides. The most prominent feature of this fold is the presence of a highly curving β hairpin formed by the last two strands of the core, which can sometimes be split into four strands. These domains do not exhibit a positively charged surface, suggesting that they might respond to ligands other than nucleic acids. (D) Representative AlphaFold models and domain schematics of Type 1D Shedu homologs with NTDs predicted to have enzymatic activity.

**Figure S8.**
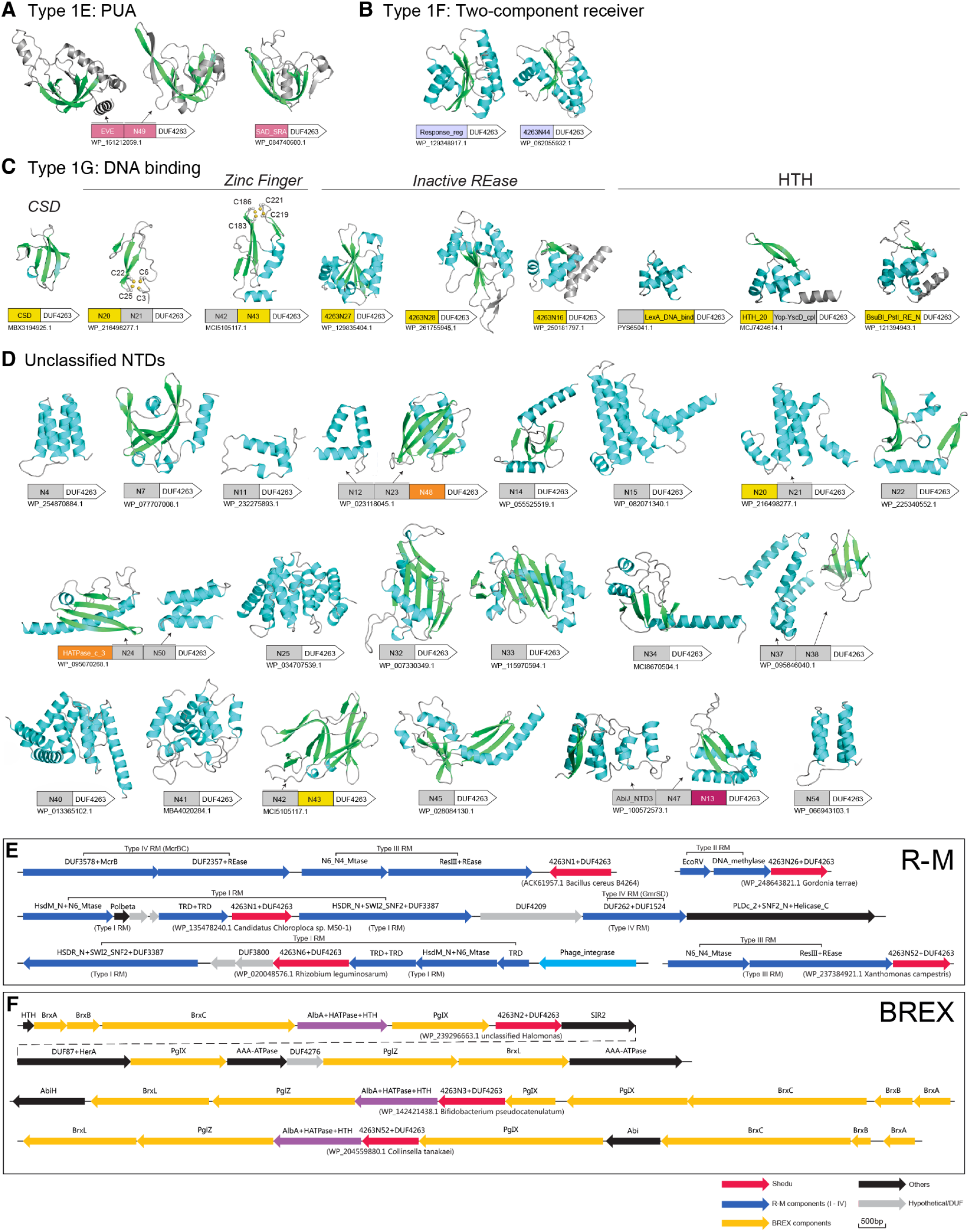
Shedu homologs encode diverse N-terminal regulatory domains (part 3) (B) Representative AlphaFold models and domain schematics of Type 1E Shedu homologs with NTDs related to the PUA fold. (C) Representative AlphaFold models and domain schematics of Type 1F Shedu homologs with NTDs related to two-component signaling systems’ phosphoreceiver domains. (D) Representative AlphaFold models and domain schematics of Type 1G Shedu homologs with NTDs predicted to bind DNA. (E) Representative AlphaFold models and domain schematics of unclassified Shedu NTDs. (F) Examples of Shedu homologs identified within restriction-modification loci. (G) Examples of Shedu homologs identified within BREX loci.

**Supplemental Table S1.**
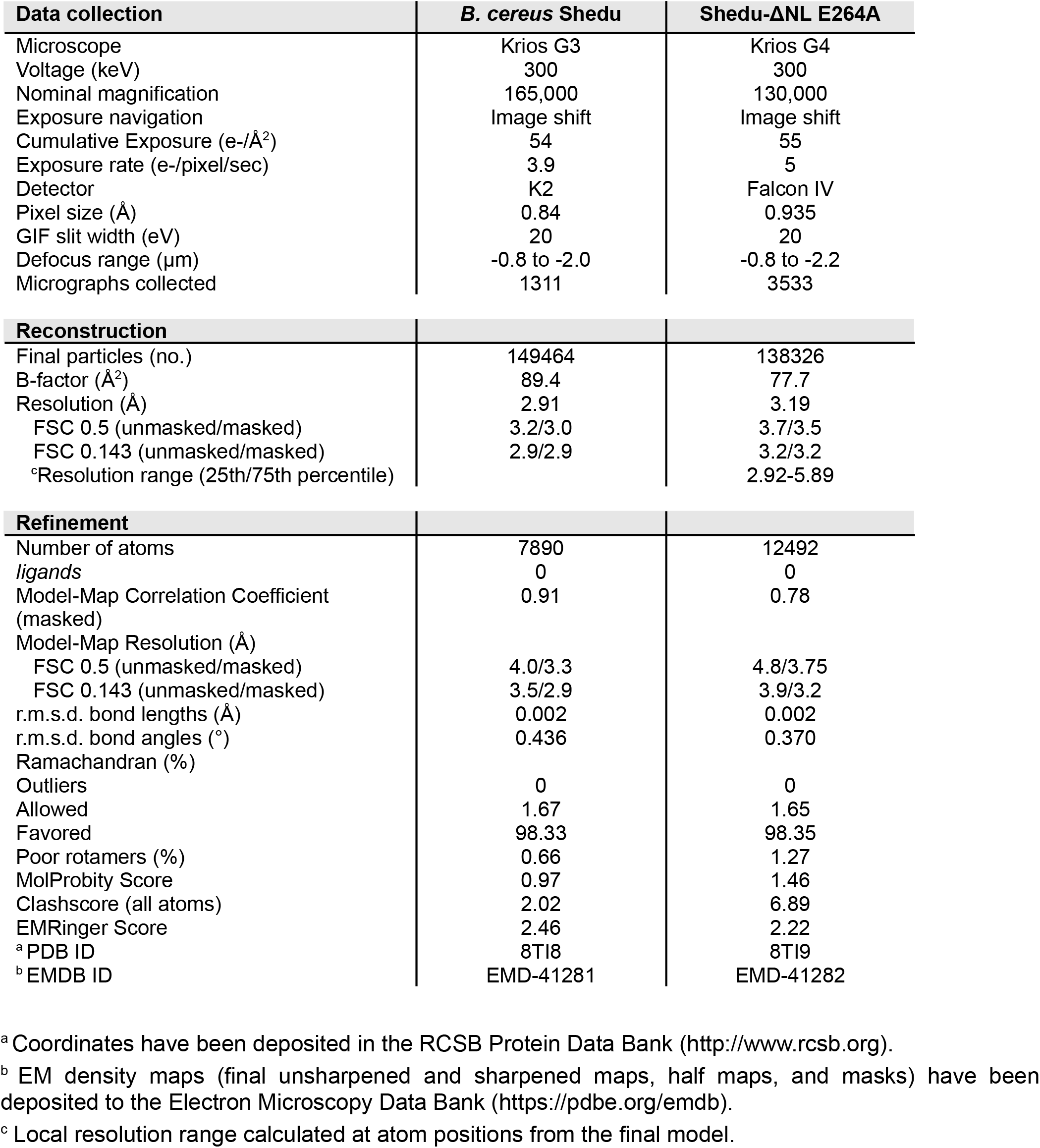
CryoEM data collection and refinement.

**Supplemental Table S2.** pUC19 sequencing after Shedu-ΔNL cleavage.

| Clone | Deletion length | Deleted bases | Full sequence length |
| --- | --- | --- | --- |
| *pUC19* | - | - | 2685 |
| 1 | 7 | 1168-1174 | 2678 |
| 2 | 7 | 323-329 | 2678 |
| 3 | 76 | 232-308 | 2609 |
| 4 | 297 | 407-703 | 2388 |
| 5 | 64 | 619-682 | 2621 |
| 6 | 237 | 207-443 | 2448 |
| 7 | 525 | 46-570 | 2160 |

**Supplemental Table S3.** List of Shedu nucleases *See attached Excel file*.

**Supplemental Movie S1. Comparison of Shedu tetramer architecture in two states**

Movie showing a morph between the active Shedu-ΔNL octamer (beginning and end of the movie) and the inactive full-length Shedu tetramer (middle of the movie). The protomer colored light blue is overlaid between the two states. View is equivalent to Figure 3C (top panel). The partially disordered linker domain and the highly mobile motif II loop are not shown.

## Notes

### Competing Interest Statement

The authors have declared no competing interest.

## References

1. Bernheim, A., and Sorek, R. (2020). The pan-immune system of bacteria: antiviral defence as a community resource. Nat. Rev. Microbiol. 18, 113–119. 10.1038/s41579-019-0278-2.

2. Labrie, S.J., Samson, J.E., and Moineau, S. (2010). Bacteriophage resistance mechanisms. Nat. Rev. Microbiol. 8, 317–327. 10.1038/nrmicro2315.

3. Tesson, F., Hervé, A., Mordret, E., Touchon, M., d’Humières, C., Cury, J., and Bernheim, A. (2022). Systematic and quantitative view of the antiviral arsenal of prokaryotes. Nat. Commun. 13, 2561. 10.1038/s41467-022-30269-9.

4. Gao, L., Altae-Tran, H., Böhning, F., Makarova, K.S., Segel, M., Schmid-Burgk, J.L., Koob, J., Wolf, Y.I., Koonin, E.V., and Zhang, F. (2020). Diverse enzymatic activities mediate antiviral immunity in prokaryotes. Science 369, 1077–1084. 10.1126/science.aba0372.

5. Doron, S., Melamed, S., Ofir, G., Leavitt, A., Lopatina, A., Keren, M., Amitai, G., and Sorek, R. (2018). Systematic discovery of antiphage defense systems in the microbial pangenome. Science 359. 10.1126/science.aar4120.

6. Payne, L.J., Meaden, S., Mestre, M.R., Palmer, C., Toro, N., Fineran, P.C., and Jackson, S.A. (2022). PADLOC: a web server for the identification of antiviral defence systems in microbial genomes. Nucleic Acids Res. 50, W541–W550. 10.1093/nar/gkac400.

7. Vassallo, C.N., Doering, C.R., Littlehale, M.L., Teodoro, G.I.C., and Laub, M.T. (2022). A functional selection reveals previously undetected anti-phage defence systems in the E. coli pangenome. Nat Microbiol 7, 1568–1579. 10.1038/s41564-022-01219-4.

8. Burroughs, A.M., Zhang, D., Schäffer, D.E., Iyer, L.M., and Aravind, L. (2015). Comparative genomic analyses reveal a vast, novel network of nucleotide-centric systems in biological conflicts, immunity and signaling. Nucleic Acids Res. 43, 10633–10654. 10.1093/nar/gkv1267.

9. Wein, T., and Sorek, R. (2022). Bacterial origins of human cell-autonomous innate immune mechanisms. Nat. Rev. Immunol. 22, 629–638. 10.1038/s41577-022-00705-4.

10. Johnson, A.G., Wein, T., Mayer, M.L., Duncan-Lowey, B., Yirmiya, E., Oppenheimer-Shaanan, Y., Amitai, G., Sorek, R., and Kranzusch, P.J. (2022). Bacterial gasdermins reveal an ancient mechanism of cell death. Science 375, 221–225. 10.1126/science.abj8432.

11. Ledvina, H.E., Ye, Q., Gu, Y., Sullivan, A.E., Quan, Y., Lau, R.K., Zhou, H., Corbett, K.D., and Whiteley, A.T. (2023). An E1–E2 fusion protein primes antiviral immune signalling in bacteria. Nature 616, 319–325. 10.1038/s41586-022-05647-4.

12. Cohen, D., Melamed, S., Millman, A., Shulman, G., Oppenheimer-Shaanan, Y., Kacen, A., Doron, S., Amitai, G., and Sorek, R. (2019). Cyclic GMP–AMP signalling protects bacteria against viral infection. Nature 574, 691–695. 10.1038/s41586-019-1605-5.

13. Hsueh, B.Y., Severin, G.B., Elg, C.A., Waldron, E.J., Kant, A., Wessel, A.J., Dover, J.A., Rhoades, C.R., Ridenhour, B.J., Parent, K.N., et al. (2022). Phage defence by deaminase-mediated depletion of deoxynucleotides in bacteria. Nat Microbiol 7, 1210–1220. 10.1038/s41564-022-01162-4.

14. Tal, N., Millman, A., Stokar-Avihail, A., Fedorenko, T., Leavitt, A., Melamed, S., Yirmiya, E., Avraham, C., Brandis, A., Mehlman, T., et al. (2022). Bacteria deplete deoxynucleotides to defend against bacteriophage infection. Nat Microbiol 7, 1200–1209. 10.1038/s41564-022-01158-0.

15. Swarts, D.C., Jore, M.M., Westra, E.R., Zhu, Y., Janssen, J.H., Snijders, A.P., Wang, Y., Patel, D.J., Berenguer, J., Brouns, S.J.J., et al. (2014). DNA-guided DNA interference by a prokaryotic Argonaute. Nature 507, 258–261. 10.1038/nature12971.

16. Cury, J., Mordret, E., Trejo, V.H., Tesson, F., Ofir, G., Poirier, E.Z., and Bernheim, A. (2022). Conservation of antiviral systems across domains of life reveals novel immune mechanisms in humans. bioRxiv, 2022.12.12.520048. 10.1101/2022.12.12.520048.

17. Morehouse, B.R., Govande, A.A., Millman, A., Keszei, A.F.A., Lowey, B., Ofir, G., Shao, S., Sorek, R., and Kranzusch, P.J. (2020). STING cyclic dinucleotide sensing originated in bacteria. Nature 586, 429–433. 10.1038/s41586-020-2719-5.

18. Bernheim, A., Millman, A., Ofir, G., Meitav, G., Avraham, C., Shomar, H., Rosenberg, M.M., Tal, N., Melamed, S., Amitai, G., et al. (2021). Prokaryotic viperins produce diverse antiviral molecules. Nature 589, 120–124. 10.1038/s41586-020-2762-2.

19. Tock, M.R., and Dryden, D.T.F. (2005). The biology of restriction and anti-restriction. Curr. Opin. Microbiol. 8, 466–472. 10.1016/j.mib.2005.06.003.

20. Roberts, R.J., Vincze, T., Posfai, J., and Macelis, D. (2015). REBASE--a database for DNA restriction and modification: enzymes, genes and genomes. Nucleic Acids Res. 43, D298–D299. 10.1093/nar/gku1046.

21. Garneau, J.E., Dupuis, M.-È., Villion, M., Romero, D.A., Barrangou, R., Boyaval, P., Fremaux, C., Horvath, P., Magadán, A.H., and Moineau, S. (2010). The CRISPR/Cas bacterial immune system cleaves bacteriophage and plasmid DNA. Nature 468, 67–71. 10.1038/nature09523.

22. Wilson, G.G., and Murray, N.E. (1991). Restriction and modification systems. Annu. Rev. Genet. 25, 585–627. 10.1146/annurev.ge.25.120191.003101.

23. Koonin, E.V., Makarova, K.S., and Zhang, F. (2017). Diversity, classification and evolution of CRISPR-Cas systems. Curr. Opin. Microbiol. 37, 67–78. 10.1016/j.mib.2017.05.008.

24. Sorek, R., Lawrence, C.M., and Wiedenheft, B. (2013). CRISPR-mediated adaptive immune systems in bacteria and archaea. Annu. Rev. Biochem. 82, 237–266. 10.1146/annurev-biochem-072911-172315.

25. Lau, R.K., Ye, Q., Birkholz, E.A., Berg, K.R., Patel, L., Mathews, I.T., Watrous, J.D., Ego, K., Whiteley, A.T., Lowey, B., et al. (2020). Structure and Mechanism of a Cyclic Trinucleotide-Activated Bacterial Endonuclease Mediating Bacteriophage Immunity. Mol. Cell 77, 723–733.e6. 10.1016/j.molcel.2019.12.010.

26. Lowey, B., Whiteley, A.T., Keszei, A.F.A., Morehouse, B.R., Mathews, I.T., Antine, S.P., Cabrera, V.J., Kashin, D., Niemann, P., Jain, M., et al. (2020). CBASS Immunity Uses CARF-Related Effectors to Sense 3’-5’- and 2’-5’-Linked Cyclic Oligonucleotide Signals and Protect Bacteria from Phage Infection. Cell 182, 38–49.e17. 10.1016/j.cell.2020.05.019.

27. Wang, S., Sun, E., Liu, Y., Yin, B., Zhang, X., Li, M., Huang, Q., Tan, C., Qian, P., Rao, V.B., et al. (2022). The complex roles of genomic DNA modifications of bacteriophage T4 in resistance to nuclease-based defense systems of E. coli. bioRxiv, 2022.06.16.496414. 10.1101/2022.06.16.496414.

28. Nakae, S., Hijikata, A., Tsuji, T., Yonezawa, K., Kouyama, K.-I., Mayanagi, K., Ishino, S., Ishino, Y., and Shirai, T. (2016). Structure of the EndoMS-DNA Complex as Mismatch Restriction Endonuclease. Structure 24, 1960–1971. 10.1016/j.str.2016.09.005.

29. Schumacher, M.A., Karamooz, E., Zíková, A., Trantírek, L., and Lukes, J. (2006). Crystal structures of T. brucei MRP1/MRP2 guide-RNA binding complex reveal RNA matchmaking mechanism. Cell 126, 701–711. 10.1016/j.cell.2006.06.047.

30. Brandsen, J., Werten, S., van der Vliet, P.C., Meisterernst, M., Kroon, J., and Gros, P. (1997). C-terminal domain of transcription cofactor PC4 reveals dimeric ssDNA binding site. Nat. Struct. Biol. 4, 900–903. 10.1038/nsb1197-900.

31. Desveaux, D., Allard, J., Brisson, N., and Sygusch, J. (2002). A new family of plant transcription factors displays a novel ssDNA-binding surface. Nat. Struct. Biol. 9, 512–517. 10.1038/nsb814.

32. Chepenik, L.G., Tretiakova, A.P., Krachmarov, C.P., Johnson, E.M., and Khalili, K. (1998). The single-stranded DNA binding protein, Pur-alpha, binds HIV-1 TAR RNA and activates HIV-1 transcription. Gene 210, 37–44. 10.1016/s0378-1119(98)00033-x.

33. Marchler-Bauer, A., Derbyshire, M.K., Gonzales, N.R., Lu, S., Chitsaz, F., Geer, L.Y., Geer, R.C., He, J., Gwadz, M., Hurwitz, D.I., et al. (2015). CDD: NCBI’s conserved domain database. Nucleic Acids Res. 43, D222–D226. 10.1093/nar/gku1221.

34. Mistry, J., Chuguransky, S., Williams, L., Qureshi, M., Salazar, G.A., Sonnhammer, E.L.L., Tosatto, S.C.E., Paladin, L., Raj, S., Richardson, L.J., et al. (2021). Pfam: The protein families database in 2021. Nucleic Acids Res. 49, D412–D419. 10.1093/nar/gkaa913.

35. Holm, L. (2022). Dali server: structural unification of protein families. Nucleic Acids Res. 50, W210–W215. 10.1093/nar/gkac387.

36. Picton, D.M., Luyten, Y.A., Morgan, R.D., Nelson, A., Smith, D.L., Dryden, D.T.F., Hinton, J.C.D., and Blower, T.R. (2021). The phage defence island of a multidrug resistant plasmid uses both BREX and type IV restriction for complementary protection from viruses. Nucleic Acids Res. 49, 11257–11273. 10.1093/nar/gkab906.

37. Machnicka, M.A., Kaminska, K.H., Dunin-Horkawicz, S., and Bujnicki, J.M. (2015). Phylogenomics and sequence-structure-function relationships in the GmrSD family of Type IV restriction enzymes. BMC Bioinformatics 16, 336. 10.1186/s12859-015-0773-z.

38. Pastor, M., Czapinska, H., Helbrecht, I., Krakowska, K., Lutz, T., Xu, S.-Y., and Bochtler, M. (2021). Crystal structures of the EVE-HNH endonuclease VcaM4I in the presence and absence of DNA. Nucleic Acids Res. 49, 1708–1723. 10.1093/nar/gkaa1218.

39. Iyer, L.M., Zhang, D., Burroughs, A.M., and Aravind, L. (2013). Computational identification of novel biochemical systems involved in oxidation, glycosylation and other complex modifications of bases in DNA. Nucleic Acids Res. 41, 7635–7655. 10.1093/nar/gkt573.

40. LeGault, K.N., Barth, Z.K., DePaola, P., and Seed, K.D. (2022). A phage parasite deploys a nicking nuclease effector to inhibit viral host replication. Nucleic Acids Res. 50, 8401–8417. 10.1093/nar/gkac002.

41. Owen, S.V., Wenner, N., Dulberger, C.L., Rodwell, E.V., Bowers-Barnard, A., Quinones-Olvera, N., Rigden, D.J., Rubin, E.J., Garner, E.C., Baym, M., et al. (2021). Prophages encode phage-defense systems with cognate self-immunity. Cell Host Microbe 29, 1620–1633.e8. 10.1016/j.chom.2021.09.002.

42. Ellis, D.M., and Dean, D.H. (1985). Nucleotide sequence of the cohesive single-stranded ends of Bacillus subtilis temperate bacteriophage phi 105. J. Virol. 55, 513–515. 10.1128/JVI.55.2.513-515.1985.

43. Perkins, J.B., Zarley, C.D., and Dean, D.H. (1978). Restriction endonuclease mapping of bacteriophage phi105 and closely related temperate Bacillus subtilis bacteriophages rho10 and rho14. J. Virol. 28, 403–407. 10.1128/JVI.28.1.403-407.1978.

44. Gao, R., and Stock, A.M. (2009). Biological insights from structures of two-component proteins. Annu. Rev. Microbiol. 63, 133–154. 10.1146/annurev.micro.091208.073214.

45. Tropea, J.E., Cherry, S., and Waugh, D.S. (2009). Expression and purification of soluble His(6)-tagged TEV protease. Methods Mol. Biol. 498, 297–307. 10.1007/978-1-59745-196-3_19.

46. Punjani, A., Rubinstein, J.L., Fleet, D.J., and Brubaker, M.A. (2017). cryoSPARC: algorithms for rapid unsupervised cryo-EM structure determination. Nat. Methods 14, 290–296. 10.1038/nmeth.4169.

47. Punjani, A., Zhang, H., and Fleet, D.J. (2020). Non-uniform refinement: adaptive regularization improves single-particle cryo-EM reconstruction. Nat. Methods 17, 1214–1221. 10.1038/s41592-020-00990-8.

48. Mirdita, M., Schütze, K., Moriwaki, Y., Heo, L., Ovchinnikov, S., and Steinegger, M. (2022). ColabFold: making protein folding accessible to all. Nat. Methods 19, 679–682. 10.1038/s41592-022-01488-1.

49. Jumper, J., Evans, R., Pritzel, A., Green, T., Figurnov, M., Ronneberger, O., Tunyasuvunakool, K., Bates, R., Žídek, A., Potapenko, A., et al. (2021). Highly accurate protein structure prediction with AlphaFold. Nature 596, 583–589. 10.1038/s41586-021-03819-2.

50. Pettersen, E.F., Goddard, T.D., Huang, C.C., Couch, G.S., Greenblatt, D.M., Meng, E.C., and Ferrin, T.E. (2004). UCSF Chimera--a visualization system for exploratory research and analysis. J. Comput. Chem. 25, 1605–1612. 10.1002/jcc.20084.

51. Emsley, P., Lohkamp, B., Scott, W.G., and Cowtan, K. (2010). Features and development of Coot. Acta Crystallogr. D Biol. Crystallogr. 66, 486–501. 10.1107/S0907444910007493.

52. Afonine, P.V., Poon, B.K., Read, R.J., Sobolev, O.V., Terwilliger, T.C., Urzhumtsev, A., and Adams, P.D. (2018). Real-space refinement in PHENIX for cryo-EM and crystallography. Acta Crystallogr D Struct Biol 74, 531–544. 10.1107/S2059798318006551.

53. Williams, C.J., Headd, J.J., Moriarty, N.W., Prisant, M.G., Videau, L.L., Deis, L.N., Verma, V., Keedy, D.A., Hintze, B.J., Chen, V.B., et al. (2018). MolProbity: More and better reference data for improved all-atom structure validation. Protein Sci. 27, 293–315. 10.1002/pro.3330.

54. Barad, B.A., Echols, N., Wang, R.Y.-R., Cheng, Y., DiMaio, F., Adams, P.D., and Fraser, J.S. (2015). EMRinger: side chain–directed model and map validation for 3D cryo-electron microscopy. Nat. Methods 12, 943–946. 10.1038/nmeth.3541.

55. Pettersen, E.F., Goddard, T.D., Huang, C.C., Meng, E.C., Couch, G.S., Croll, T.I., Morris, J.H., and Ferrin, T.E. (2021). UCSF ChimeraX: Structure visualization for researchers, educators, and developers. Protein Sci. 30, 70–82. 10.1002/pro.3943.

56. Kropinski, A.M., Mazzocco, A., Waddell, T.E., Lingohr, E., and Johnson, R.P. (2009). Enumeration of bacteriophages by double agar overlay plaque assay. Methods Mol. Biol. 501, 69–76. 10.1007/978-1-60327-164-6_7.

57. Altschul, S.F., Madden, T.L., Schäffer, A.A., Zhang, J., Zhang, Z., Miller, W., and Lipman, D.J. (1997). Gapped BLAST and PSI-BLAST: a new generation of protein database search programs. Nucleic Acids Res. 25, 3389–3402. 10.1093/nar/25.17.3389.

58. Fu, L., Niu, B., Zhu, Z., Wu, S., and Li, W. (2012). CD-HIT: accelerated for clustering the next-generation sequencing data. Bioinformatics 28, 3150–3152. 10.1093/bioinformatics/bts565.

59. Frickey, T., and Lupas, A. (2004). CLANS: a Java application for visualizing protein families based on pairwise similarity. Bioinformatics 20, 3702–3704. 10.1093/bioinformatics/bth444.

60. Eddy, S.R. (2011). Accelerated Profile HMM Searches. PLoS Comput. Biol. 7, e1002195. 10.1371/journal.pcbi.1002195.

61. Pei, J., Kim, B.-H., and Grishin, N.V. (2008). PROMALS3D: a tool for multiple protein sequence and structure alignments. Nucleic Acids Res. 36, 2295–2300. 10.1093/nar/gkn072.

62. Holm, L., and Park, J. (2000). DaliLite workbench for protein structure comparison. Bioinformatics 16, 566–567. 10.1093/bioinformatics/16.6.566.

63. Holm, L., and Laakso, L.M. (2016). Dali server update. Nucleic Acids Res. 44, W351–W355. 10.1093/nar/gkw357.

64. Jurrus, E., Engel, D., Star, K., Monson, K., Brandi, J., Felberg, L.E., Brookes, D.H., Wilson, L., Chen, J., Liles, K., et al. (2018). Improvements to the APBS biomolecular solvation software suite. Protein Sci. 27, 112–128. 10.1002/pro.3280.

65. Schneider, T., Tan, Y., Li, H., Fisher, J.S., and Zhang, D. (2022). Photoglobin, a distinct family of non-heme binding globins, defines a potential photosensor in prokaryotic signal transduction systems. Comput. Struct. Biotechnol. J. 20, 261–273. 10.1016/j.csbj.2021.12.022.

